# SDS-22 stabilizes the PP1 catalytic subunits GSP-1/-2 contributing to polarity establishment in *C. elegans* embryos

**DOI:** 10.1101/2025.01.07.631699

**Authors:** Yi Li, Ida Calvi, Monica Gotta

## Abstract

In many cells, cell polarity depends on the asymmetric distribution of the conserved PAR proteins, maintained by a balanced activity between kinases and phosphatases. The *C. elegans* one-cell embryo is polarized along the anterior-posterior axis, with the atypical protein kinase C PKC-3 enriched in the anterior, and the ring finger protein PAR-2 in the posterior. PAR-2 localization is regulated by PKC-3 and the PP1 phosphatases GSP-1/-2. Here, we find that, similar to GSP-2 depletion, depletion of the conserved PP1 interactor SDS-22 results in a partial rescue of the polarity defects of a *pkc-3* temperature-sensitive mutant. Consistent with the rescue, SDS-22 depletion or mutation results in reduced GSP*-*1/*-*2 protein levels and activity. The decreased levels of GSP-1/-2 can be rescued by reducing proteasomal activity. Our data suggest that SDS-22 contributes to polarity by protecting the GSP-1 and GSP-2 catalytic subunits from proteasome-mediated degradation, supporting recent data in human cells showing the SDS22 is required to stabilize nascent PP1.

## Introduction

Cell polarity is crucial for cellular and tissue architecture and function. In epithelial cells, polarity specifies the apical, basal and lateral membrane domains, enabling the selective barrier function of the epithelium (Riga *et al*, 2020). In migrating cells, front-to-rear polarity is crucial for cells to respond to environmental cues such as inflammatory factors, bacterial products, or growth factors (Llense & Etienne-Manneville, 2015). Cell polarity is crucial for asymmetric cell division, which is a prerequisite for stem cells to self-renew and to generate specialized daughter cells (Santoro *et al*, 2016) and for embryos to properly develop (Campanale *et al*, 2017). In many cells, polarity is regulated by the conserved partitioning defective (PAR) proteins, which were first identified in *C. elegans* (Goldstein & Macara, 2007; Lang & Munro, 2017).

The one-cell *C. elegans* embryo is polarized along the anterior-posterior axis. The anterior PAR proteins (the PDZ proteins PAR-3 and PAR-6, the atypical protein kinase C PKC-3 and the small GTPases CDC-42) are localized to the anterior cortex and cytoplasm of the embryo, while the posterior PAR proteins (the kinase PAR-1, the ring finger protein PAR-2, the lethal giant larvae ortholog LGL-1 and the CDC-42 GAP CHIN-1) are enriched in the posterior half of the embryo (reviewed in (Goehring, 2014; Lang & Munro, 2017)). Before polarity establishment, the anterior PAR proteins are uniformly localized on the cortex, while the posterior PAR proteins are localized in the cytoplasm. Shortly after fertilization, the centrosomal mitotic kinase Aurora A (AIR-1) removes the Rho GEF ECT-2 from the posterior pole (Kapoor & Kotak, 2019; Klinkert *et al*, 2019; Longhini & Glotzer, 2022; Zhao *et al*, 2019). This event initiates a cortical flow that segregates the anterior PAR proteins to the anterior and liberates the posterior cortex of the embryo, which is then occupied by the posterior PAR proteins (Munro *et al*, 2004).

Several studies have emphasized the importance of the regulation of PAR-2 phosphorylation by PKC*-*3 in establishing and maintaining cortical polarity (Hao *et al*, 2006; Motegi *et al*, 2011). Recently, our group has shown that PP1 phosphatases are also required for cortical polarity in one-cell embryos (Calvi *et al*, 2022). Before polarity establishment, PKC-3 phosphorylates PAR-2 and inhibits PAR-2 localization at the cortex (Hao *et al*., 2006; Motegi *et al*., 2011). The PP1 phosphatases GSP-1/-2 dephosphorylate PAR-2 allowing its cortical posterior accumulation and embryo polarization. Phosphomimetic mutants of PAR-2 (Hao *et al*., 2006), or mutations that abolish the interaction between PAR-2 and GSP-1/-2 (Calvi *et al*., 2022), prevent PAR-2 localization at the posterior, indicating that PP1 phosphatases are crucial for polarity establishment in *C. elegans* embryos.

Analogous to the intricate regulation of protein kinases, the activity of phosphatases towards specific substrates must be tightly regulated for proper cellular function, including cell division, heat shock response, and glycogen metabolism (Ceulemans & Bollen, 2004). The activity of PP1 is primarily regulated post-translationally through interactions with PP1 interacting proteins, which guide its localization, substrate specificity and function. Over 200 PP1 interactors have been reported in different species and they regulate both in time and space the targeting of catalytic subunits towards different substrates (Aggen *et al*, 2000; Verbinnen *et al*, 2017; Virshup & Shenolikar, 2009). One non-canonical regulator of PP1 is the conserved SDS22 protein. In mammalian cells SDS22 suppresses Aurora B activity to control cell division (Duan *et al*, 2016; Posch *et al*, 2010; Wurzenberger *et al*, 2012). In *Drosophila melanogaster*, kinetochore-localized PP1–Sds22 dephosphorylates moesin at cell poles, facilitating anaphase elongation through polar relaxation (Rodrigues *et al*, 2015). PP1-87B and Sds22 also counteract aPKC and Aurora A kinase phosphorylation of Lgl, regulating apical-basal polarity in follicle epithelial cells (Moreira *et al*, 2019). In *Saccharomyces cerevisiae*, Sds22, along with Ypi1, is critical for the nuclear import and stabilization of PP1-Glc7 (Cheng & Chen, 2015; Peggie *et al*, 2002). Depletion of either Sds22 or Ypi1 results in mitotic arrest (Pedelini *et al*, 2007). Recent biochemical studies (Kueck *et al*, 2024) and studies in human cells (Cao *et al*, 2024) have demonstrated that SDS22 stabilizes nascent PP1 but needs to be removed from PP1 for the enzyme to be active. These two recent findings suggest that while SDS-22 is required for the biogenesis of PP1 holoenzymes, its removal is essential to have an active PP1. This dual role of SDS-22 explains how SDS22 behaves as an inhibitor in biochemical assays in vitro but as an activator in vivo (Cao *et al*., 2024; Cao *et al*, 2022; Kueck *et al*., 2024; Lesage *et al*, 2007).

In *C. elegans* GSP-1 and GSP-2, orthologues of human PP1β and PP1α, contribute to diverse processes such as germline immortality (Billmyre *et al*, 2019), sister chromatid cohesion in meiosis (Hsu *et al*, 2000; Tzur *et al*, 2012), centriole duplication (Peel *et al*, 2017), PAR-2 localization (Calvi *et al*., 2022) in the early embryos, and the maintenance of epidermal junctions in adult worms (Beacham *et al*, 2022). Associated regulatory proteins have been identified for specific functions: for example, SDS-22 and SZY-2 modulate GSP-1 and GSP-2 activity in centriole duplication (Peel *et al*., 2017), and APE-1 directs localization of GSP-2 to epidermal cell-cell junctions (Beacham *et al*., 2022). However, the regulation of GSP-1 and GSP-2 during polarity establishment in one-cell embryos remains unknown.

Here we show that SDS-22 depletion partially rescues the polarity defects caused by reduced PAR-2 phosphorylation in the *pkc-3(ne4246)* mutant at the semi-restrictive temperature (24°C). Depletion of SDS-22 results in lower GSP-1 and GSP-2 protein levels which can be rescued by depleting proteasomal subunits. These results establish that SDS-22 contributes to cell polarity by regulating GSP-1/-2 levels and are consistent with and complement the recent data in mammalian cells showing that SDS22 is important to control the stability of the PP1 phosphatase (Cao *et al*., 2024).

## Result

### Depletion of SDS-22 suppresses polarity defects of a temperature sensitive *pkc-3* mutant

We have previously found that PP1 phosphatases GSP-1 and GSP-2 counteract the activity of the atypical protein kinase PKC-3 and allow cortical localization of PAR-2 in *C. elegans* embryos. However, the regulator(s) of GSP-1 and GSP-2 in polarity establishment is not known. In immunoprecipitations followed by mass spectrometry analysis of mNG::GSP-2 from embryos, we identified three interactors that have been previously reported as evolutionarily conserved PP1 regulators: SDS-22, APE-1 and SYZ-2(I-2) (Fig EV1, Dataset EV1 and EV2).

When the temperature sensitive mutant *pkc-3(ne4246)* is grown at semi-permissive temperature, the residual PKC-3 activity is not sufficient to exclude PAR-2 from the anterior cortex. These embryos cannot establish polarity and die. Depletion of GSP-2 in this strain suppresses PAR-2 mislocalization and the resulting polarity defects, thereby rescuing embryonic lethality. We first asked whether depletion of any of these three identified regulators suppresses the embryonic lethality of *pkc-3(ne4246); gfp::par-2* embryos at the semi-permissive temperature of 24°C (temperature used in all experiments with the *pkc-3(ne4246)* mutant, unless otherwise stated), similar to depletion of GSP-2. None of these three regulators was able to suppress the embryonic lethality (Fig 1A). SDS-22 depletion alone resulted in 73.6% of embryonic lethality in the *gfp::par-2* strain, consistent with previous findings (Peel *et al*., 2017). We then investigated if any of these regulators was able to rescue the aberrant PAR-2 localization of *pkc-3(ne4246)* embryos. Neither APE-1 nor SYZ-2 depletion rescued the symmetric PAR-2 localization observed in *pkc-3(ne4246); gfp::par-2* embryos (Fig EV2A). In contrast, depletion of SDS-22 resulted in PAR-2 localization being enriched in the posterior cortex in 87.5% of the one-cell stage embryos (Fig 1B,C) and PAR-2 was localized to the P1 blastomere after the first cell-division (Movie EV1). Depletion of SDS-22 in otherwise wildtype *gfp::par-2* embryos did not impair anterior-posterior polarity, with PAR-2 restricted to the posterior cortex (Fig 1D), indicating that the lethality caused by loss of SDS-22 is not the result of impaired PAR polarity establishment. However, similar to what has been observed in GSP-2 depleted embryos (Calvi *et al*., 2022), the PAR-2 domain was smaller (Fig 1D, Fig EV2B). SDS-22 depleted embryos also displayed a smaller size (Fig EV2C).

**Fig 1:**
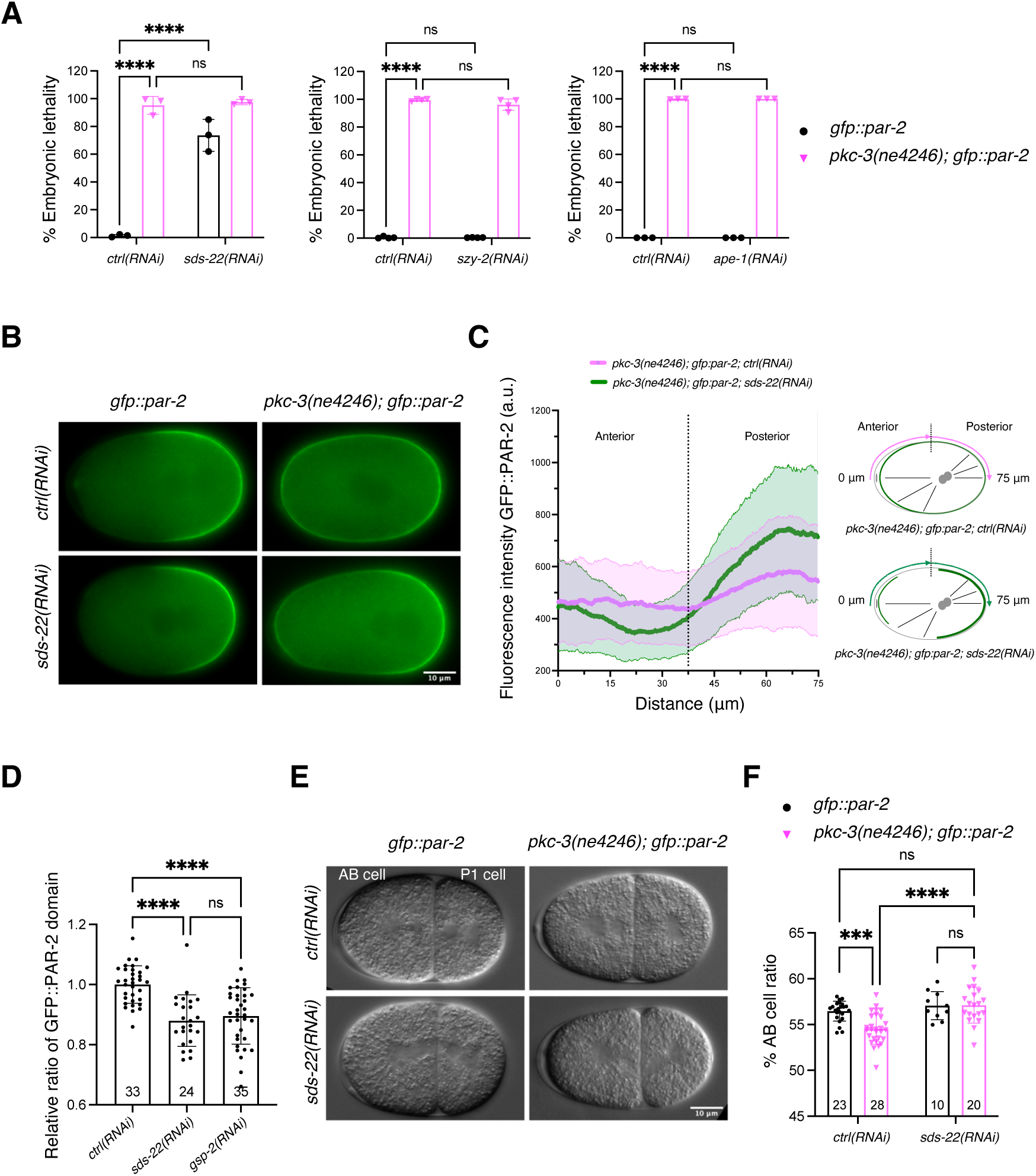
SDS-22 depletion partially suppresses the polarity defects of the *pkc-3(ne4246)* mutant. **(A)** Embryonic lethality of *gfp::par-2* and *pkc-3(ne4246); gfp::par-2* embryos with the indicated depletions. The reported values correspond to the percentage of unhatched embryos over the total progeny (larvae and unhatched embryos) in all figures. *gfp::par-2; ctrl(RNAi)*, *n* = 4,975, *gfp::par-2; sds-22(RNAi)*, *n* = 3,923, *pkc-3(ne4246); gfp::par-2; ctrl(RNAi)*, *n* = 2,247, and *pkc-3(ne4246); gfp::par-2; sds-22(RNAi)*, *n* = 1,611. *gfp::par-2; szy-2(RNAi)*, *n* = 3,550; *pkc-3(ne4246); gfp::par-2; szy-2(RNAi)*, *n* = 2,588. *gfp::par-2; ape-1(RNAi)*, *n* = 720; *pkc-3(ne4246); gfp::par-2; ape-1(RNAi)*, *n* = 930. *sds-22(RNAi)* and *ape-1(RNAi), N* = 3. *szy-2(RNAi), N* = 4. The mean is shown and error bars indicate Standard Deviation (SD). The P values were determined using two-way ANOVA Tukey’s multiple comparisons test. For plots of embryonic lethality, each dot inside the bar represents the average lethality of each independent experiment, unless otherwise indicated. **(B)** Representative frames of time-lapse videos of embryos at the pronuclear meeting stage: *gfp::par-*2; *ctrl(RNAi)*, *n* = 12 (PAR-2 is posterior in 100% of them), *gfp::par-2; sds-22(RNAi), n* = 12 (100% posterior PAR-2), *pkc-3(ne4246); gfp::par-2; ctrl(RNAi), n* = 42 (0% posterior PAR-2, PAR-2 is all around the cortex in all of them), and *pkc-3(ne4246); gfp::par-2; sds-22(RNAi*), *n* = 48 (87.5% have PAR-2 enriched in the posterior cortex)*. N* = 5. **(C)** GFP::PAR-2 intensity line profile along half of the embryo cortex starting from the anterior, as shown in the scheme on the right. *pkc-3(ne4246); gfp::par-2; ctrl(RNAi), n* = 31, *pkc-3(ne4246); gfp::par-2; sds-22(RNAi), n* = 32*. N* = 5. **(D)** Quantification of the GFP::PAR-2 domain size at pronuclear meeting in live embryos with the indicated RNAi conditions. Sample size (*n*) is indicated inside the bars in the graph. *N* = 4. Mean is shown and error bars indicate SD. The P values were determined using one-way ANOVA. In C and E, each dot represents a single embryo. **(E)** Still images of two-cell embryos of the indicated genotypes taken from DIC time-lapse movies. **(F)** Quantification of the AB cell size as a percentage of the whole embryo size. Sample size (*n*) is indicated inside the bars in the graph. *N* = 3. Mean is shown and error bars indicate SD. The P-values were determined using two-way ANOVA “Tukey’s multiple comparisons test”. For all the panels, RNA interference was performed by feeding. Scale bar is 10 µm, anterior is to the left and posterior to the right. In all plots, ns p > 0.05, * p < 0.05, ** p < 0.01, ***p < 0.001, ****p < 0.0001, *n* = number of embryos analyzed; *N* = number of independent experiments.

We then investigated whether SDS-22 depletion, similar to GSP-2 depletion, rescued aberrant PAR-2 localization in another genetic background. When the PP1 binding motif in PAR-2 (164 RLFF 167) is optimized by a single amino acid substitution (L165V, resulting in RVFF, based on previous studies that show better binding to PP1(Meiselbach *et al*, 2006)), PAR-2 abnormally occupies the anterior cortex in early embryos. This phenotype is rescued by depletion of GSP-2, suggesting that it depends on higher activity of GSP-2 on PAR-2 (Calvi *et al*., 2022). SDS-22 depletion also rescued the anterior PAR-2 domain observed in *gfp::par-2(L165V)* embryos as anterior PAR-2 was observed in only 45.5% of *gfp::par-2(L165V); sds-22(RNAi*) embryos compared to 100% *gfp::par-2(L165V); ctrl(RNAi*) embryos (Fig EV3).

Polarity controls the posterior positioning of the mitotic spindle, which results in an asymmetric cell division producing a larger anterior AB cell and a smaller posterior P1 cell. In *gfp::par-2* embryos AB cell accounts for 56.5% of the total embryo length. In contrast, polarity defects in the *pkc-3(ne4246)* mutant lead to reduced cell size asymmetry with a smaller AB cell, occupying 54.2% of the embryo length. Consistent with the suppression of PAR-2 polarity defect, depletion of SDS-22 in *pkc-3(ne4246)* embryos resulted in a bigger AB cell (57.1% embryo length), comparable to the size of AB in control *gfp::par-2* embryos (Fig 1E,F).

Our data show that depletion of SDS-22 results in a smaller PAR-2 domain, partially suppresses the polarity defects of a *pkc-3* temperature sensitive strain and the aberrant PAR-2 localization observed in the PAR-2(L165V) mutant strain. As SDS-22 is a conserved PP1 regulator, our data suggest that SDS-22 positively regulates GSP-2.

### The conserved E153 of SDS-22 is important for SDS-22 interaction with GSP-1 and GSP-2

We have identified SDS-22 in immunoprecipitation with GSP-2 from embryos, thereby confirming a previous finding that SDS-22 immunoprecipitates with both GSP-1 and GSP-2 from full worm lysates (Peel *et al*., 2017). We were able to confirm the interaction of SDS-22 with both GSP-1 and GSP-2 in a two-hybrid assay (Fig 2).

**Fig 2:**
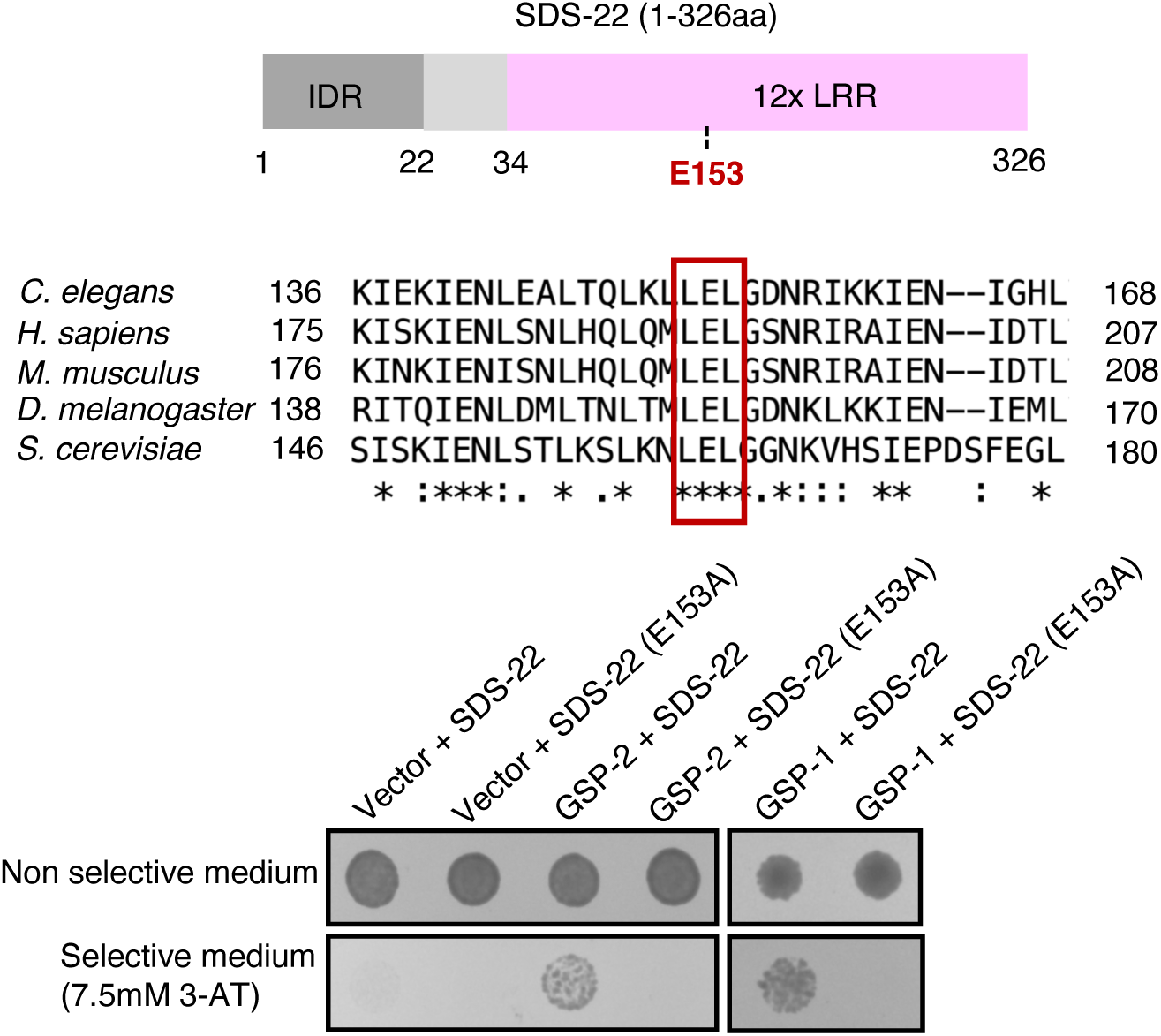
SDS-22 interacts with GSP-1 and GSP-2 via the conserved E153 residue. Upper panel: schematic representation of the full-length (1-326aa) SDS-22. IDR: intrinsically disordered region; LRR: leucine-rich repeats; E153: glutamic acid at amino acid residue 153. Middle panel: sequence alignment of one PP1 interacting region in SDS22 from *C. elegans* (Uniprot P45969), *H. sapiens* (Uniprot Q15435), *M. musculus* (Uniprot Q3UM45), *D. melanogaster* (Uniprot Q9VEK8), and *S. cerevisiae* (Uniprot 936047). The reported PP1-binding residue E192 of *H. sapiens* is present in *C. elegans* as E153, and this site is conserved in all species listed (marked by red square). Figure adapted from alignment obtained with Uniprot alignment tool (https://www.uniprot.org/align). Lower panel: yeast two-hybrid assays showing the interaction between SDS-22 (wild type and the E153A mutant) and GSP-1 and GSP-2. Yeasts were transformed with the indicated plasmids (SDS-22 and SDS-22(E153A) are fused to the DNA Binding Domain and GSP-1 and GSP-2 to the Activation Domain) and were grown on non-selective and selective medium. Interaction between bait and prey results in growth on selective medium (7.5mM 3-AT).

SDS-22 is a leucine-rich repeat (LRR) protein and does not contain the degenerate RLxF PP1 docking motif that has been found in most PP1 regulators. Instead, a conserved glutamic acid E153 in the LRR domain is important for its interaction with the PP1 catalytic subunits (Fig 2) (Ceulemans *et al*, 2002; Eiteneuer *et al*, 2014; Heroes *et al*, 2019). This site is conserved in *C. elegans* and we asked whether it is important for the interaction of SDS-22 with GSP-1 and GSP-2. Substitution of the glutamine E153 with Alanine (referred to as SDS-22(E153A)) abrogated the interaction with both GSP-1 and GSP-2 in the two-hybrid assays (Fig 2).

These data confirm that SDS-22 interacts with both GSP-1 and GSP-2 and identify the glutamic acid residue E153 in the LRR domain of SDS-22 as crucial for the interaction, consistent with what has been demonstrated for the human orthologs (Ceulemans *et al*., 2002; Eiteneuer *et al*., 2014; Heroes *et al*., 2019).

### Polarity defects are rescued in pkc-3(ne4246); sds-22(E153A); gfp::par-2 embryos

Our two-hybrid data show that SDS-22 interacts with GSP-1 and GSP-2 through the E153 residue. We introduced this substitution in vivo and asked whether *sds-22(E153A)* mutant embryos display a smaller PAR-2 domain, as observed in SDS-22 depleted embryos (Fig 1B,D). PAR-2 was restricted to the posterior in *gfp::par-2; sds-22(E153A)* as in control embryos, but the length of the domain was decreased (Fig 3A,B, Movie EV2), similar to what was observed in SDS-22 and GSP-2 depleted embryos. However, while SDS-22 depletion resulted in 73.6% embryonic lethality (Fig 1A), *gfp::par-2; sds-22(E153A)* only exhibited 5.3% embryonic lethality compared to the *gfp::par-2* strain (Fig 3C). Because of this, we decided to test whether *sds-22(E153A)* was able to rescue the lethality and the polarity defects of *pkc-3(ne4246)* embryos. We introduced the E153A mutation in *pkc-3(ne4246); gfp::par-2* strain, let them grow at the semi-permissive temperature (24°C) and quantified embryonic lethality. We found that the lethality of the embryos was not rescued at this temperature (Fig 3D).

**Fig 3:**
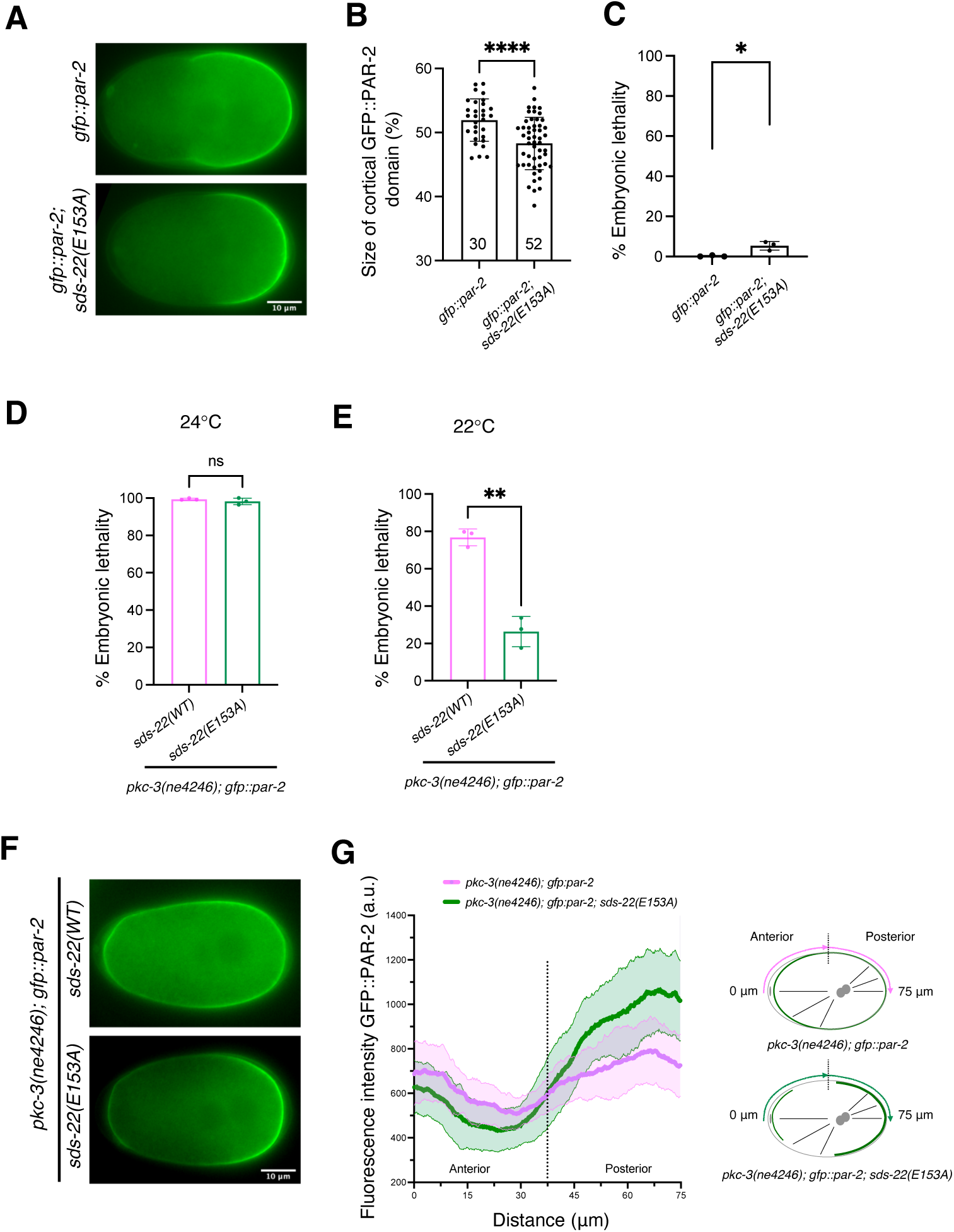
Mutation of SDS-22 E153 residue partially rescues polarity defects of *pkc-3(ne4246)* mutant. **(A)** Still images at the pronuclear meeting stage from time-lapse videos of *gfp::par-2* and *gfp::par-2; sds-22(E153A)*. **(B)** Quantification of the GFP::PAR-2 domain size at pronuclear meeting in live zygotes. Sample size (*n*) is indicated inside the bars in the graph, each dot represents a single embryo. *N* = 3. Mean is shown and error bars indicate SD. The P value was determined using two-tailed unpaired Student’s *t* test. **(C)** Percentage of embryonic lethality of *gfp::par-2* and *gfp::par-2; sds-22(E153A). gfp::par-2, n* = 757, *gfp::par-2; sds-22(E153A), n* = 2644. *N* = 3. Mean is shown and error bars indicate SD. The P value was determined using two-tailed unpaired Student’s *t* test. **(D, E)** Percentage of embryonic lethality of *pkc-3(ne4246); gfp::par-2* and *pkc-3(ne4246); gfp::par-2; sds-22(E153A)* at the semi-permissive temperature of 24°C **(D)** and of 22°C **(E)**. For **(D)**, *pkc-3(ne4246); gfp::par-2*, *n* = 569, and *pkc-3(ne4246); gfp::par-2; sds-22(E153A), n* = 545. For **(E)**, *pkc-3(ne4246); gfp::par-2*, *n* = 2171, and *pkc-3(ne4246); gfp::par-2; sds-22(E153A) n* = 1867. *N* = 3. Mean is shown and error bars indicate SD. The P value was determined using two-tailed unpaired Student’s *t* test. In C, D and E, dots represent the mean lethality of each independent experiment. **(F)** Still images from time-lapse videos of *pkc-3(ne4246); gfp::par-2* and *pkc-3(ne4246); gfp::par-2; sds-22(E153A)* embryos at the pronuclear meeting stage. Worms were grown at the semi-permissive temperature of 22°C as in panel E. **(G)** GFP::PAR-2 intensity line profile along half of the embryo cortex starting from the anterior, as shown in the scheme on the right. *pkc-3(ne4246); gfp::par-2* and *pkc-3(ne4246); gfp::par-2; sds-22(E153A) n* = 19 for both genotypes*. N* = 3. Worms were grown at the semi-permissive temperature of 22°C as in panel E and F. Scale bar is 10 µm, anterior is to the left and posterior to the right. In all plots, ns p > 0.05, * p < 0.05, ** p < 0.01, ***p < 0.001, ****p < 0.0001, *n* = number of embryos analyzed; *N* = number of independent experiments.

Decreasing the temperature to 22°C reduced the embryonic lethality in the *pkc-3(ne4246); gfp::par-2* strain (Fig 3E) to about 76.8%. In these more permissive conditions, *pkc-3(ne4246); sds-22(E153A); gfp::par-2* exhibited only 26.4% embryonic lethality, indicating that the *sds-22(E153A)* mutation, which abolishes SDS-22 interaction with GSP-1/-2, can partially rescue the lethality of *pkc-3(ne4246)* at this temperature.

When we looked at polarity we found that PAR-2, despite still weakly detectable at the anterior, was more enriched at the posterior cortex in the *pkc-3(ne4246); sds-22(E153A); gfp::par-2* mutant embryos compared to *pkc-3(ne4246); gfp::par-2* (Fig 3F,G, Movie EV 3). The rescue of embryonic lethality and polarity phenotypes was not due to a reduced expression level of SDS-22(E153A) because SDS-22(E153A)::GFP showed similar protein intensity as SDS-22::GFP (Fig EV4).

Our data show that a mutation that disrupts the interaction of SDS-22 with GSP-1 and GSP-2 in a two-hybrid assay, can lead to a partial rescue of the lethality and the polarity defect of *pkc-3(ne4246); gfp::par-2* embryos, suggesting that the interaction of SDS-22 with the PP1 phosphatases contributes to polarity establishment.

### Loss or mutation of SDS-22 results in hyper-phosphorylation of a PP1 substrate

In *C. elegans* embryos GSP-1 and GSP-2 redundantly regulate PAR-2 localization (Calvi *et al*., 2022). GSP-2 is the predominant PP1 phosphatase that allows for PAR-2’s localization to the posterior cortex. Depletion of GSP-2 reduces the size of the PAR-2 domain in the *gfp::par-2* strain while depletion of GSP-1 does not lead to observable defects in PAR-2 domain size. Only when GSP-1 and GSP-2 are co-depleted, PAR-2 remains mostly cytoplasmic (Fig 4A, (Calvi *et al*., 2022)). If SDS-22 was essential to regulate the activity of both GSP-1 and GSP-2, its depletion should result in a loss of PAR-2 cortical localization, similar to what is observed in the co-depletion of GSP-1 and GSP-2. As shown above, depletion of SDS-22 in the *gfp::par-2* strain did not lead to cytoplasmic localization of PAR-2 (Fig 1B and 4A) but resulted in a smaller PAR-2 domain (Fig 1D and 4B), as observed in the GSP-2 depletion. These data suggest that SDS-22 regulates GSP-2 rather than GSP-1. If this was true, depletion of both SDS-22 and GSP-1 should result in PAR-2 remaining in the cytoplasm, similar to GSP-1 and GSP-2 co-depletion. We therefore co-depleted SDS-22 and GSP-1 and, as control, SDS-22 and GSP-2. The double depletion was efficient as shown in Fig EV5. In embryos co-depleted of SDS-22 and GSP-1, PAR-2 still formed a posterior cortical domain. Co-depletion of SDS-22 and GSP-2 resulted in an even more pronounced reduction of PAR-2 domain compared to the single depletion of SDS-22 (Fig 4A,B), suggesting that SDS-22 plays a different role than simply activating GSP-2 (see below).

**Fig 4:**
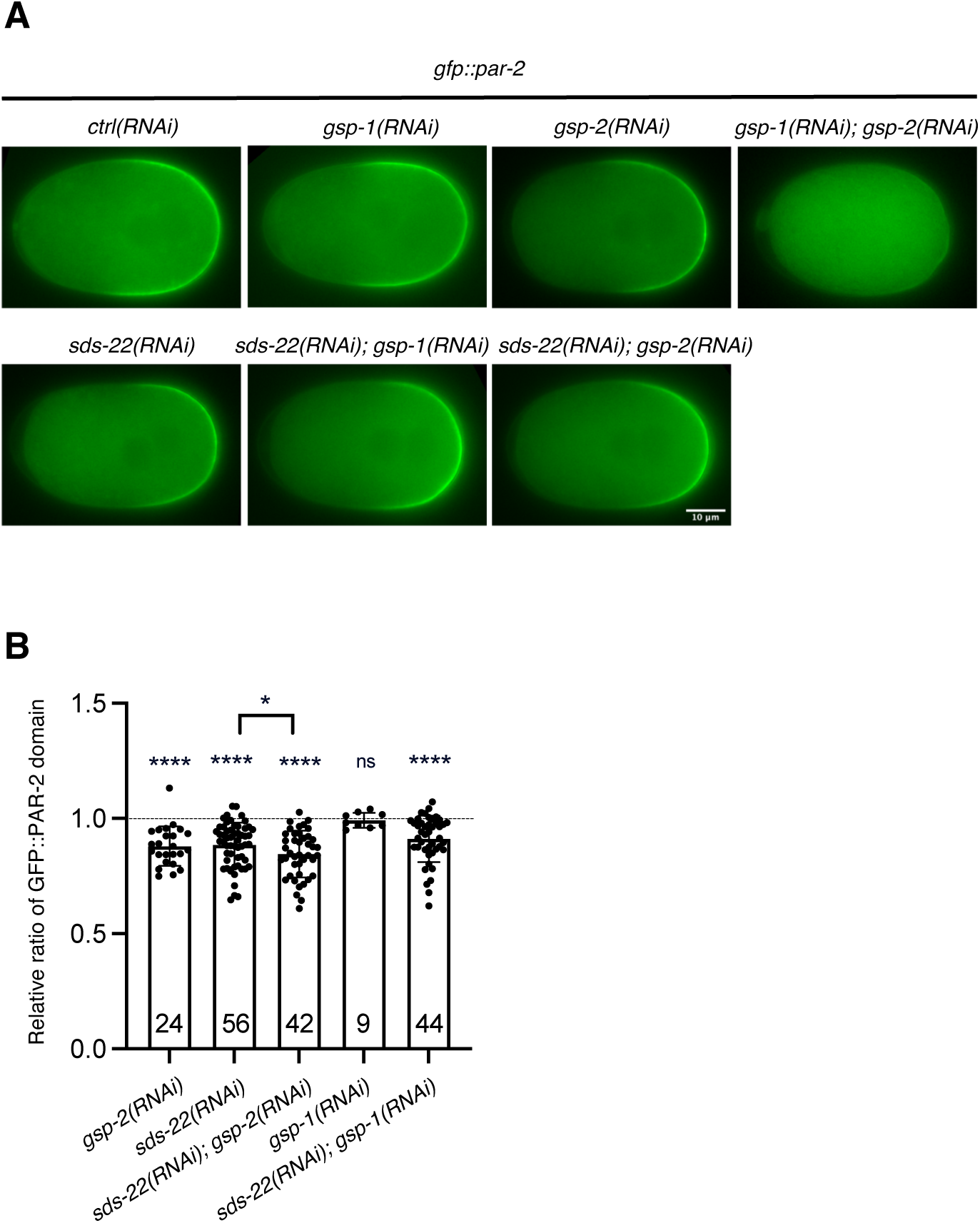
SDS-22 does not genetically regulate GSP-1, GSP-2 or both. **(A)** Still images from time-lapse videos of *gfp::par-2* zygotes at pronuclear meeting with the indicated depletions. RNA interference was performed by injection. Scale bars is 10 µm, anterior is to the left and posterior to the right. **(B)** Quantification of the relative ratio of GFP::PAR-2 size domain with indicated RNAi conditions. Each condition was compared to *ctrl(RNAi)* treatment within the same experiment. The P value was determined using two-tailed unpaired Student’s *t* test. Sample size (*n*) is indicated inside the bars in the graph. *N* = 4-5. Mean is shown and error bars indicate SD. ns p > 0.05, * p < 0.05, ** p < 0.01, ***p < 0.001, ****p < 0.0001.

Our findings show that depletion or mutation of SDS-22 partially rescues polarity defects of the *pkc-3(ne4246)* mutant. However, neither SDS-22 depletion alone nor co-depletion with GSP-1 or GSP-2 prevents PAR-2 cortical localization similar to GSP-1/-2 co-depletion. To understand the role of SDS-22 in the regulation of GSP-1 and GSP-2, we investigated whether SDS-22 depletion or mutation decreases PP1 activity. To quantify the activity of the PP1 phosphatase, we examined the phosphorylation levels of a well characterized substrate, histone H3, as previously described (Hsu *et al*., 2000; Qian *et al*, 2011). Histone H3 is phosphorylated at the Ser 10 residue by the Aurora B kinase in the early phases of mitosis and it is dephosphorylated by PP1 at anaphase (Xin *et al*, 2020). We found that phospho-histone H3 (Ser10) levels on the chromosomes during anaphase were higher in SDS-22 depleted one-cell embryos (Fig 5A) compared to control depletion, indicating that depletion of SDS-22 leads to deficient dephosphorylation of PP1 substrates.

**Fig 5.**
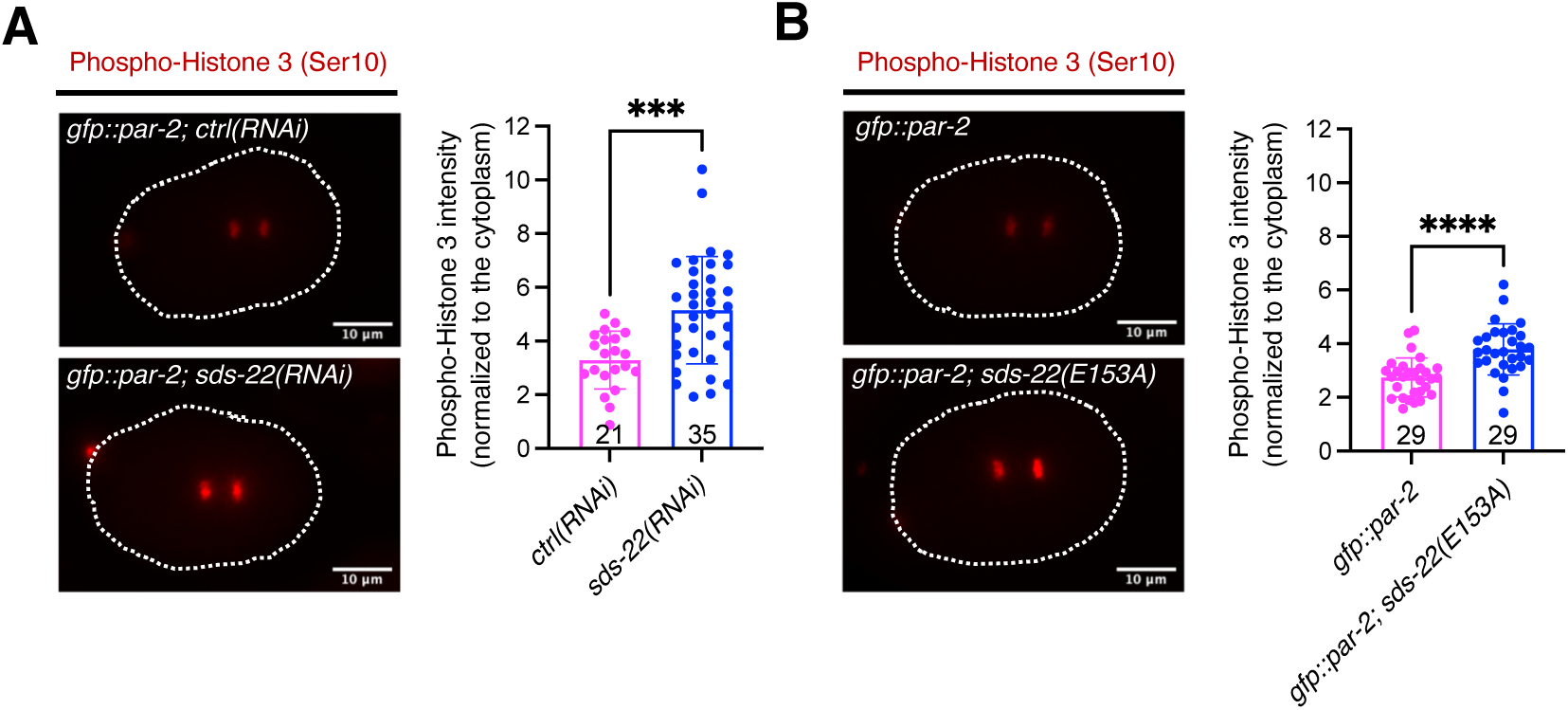
Loss or mutation of SDS-22 decreases GSP-1 and GSP-2 activity. **(A)** Left, images of a *gfp::par-2; ctrl(RNAi)* anaphase embryo and a *gfp::par-2; sds-22(RNAi)* anaphase embryo stained for phospho-histone H3 (Ser10) (in red). Right, quantifications. **(B)** Left, images of a *gfp::par-2* and a *gfp::par-2*; *sds-22(E153A)* embryo at anaphase stained for phospho-histone H3(Ser10) (in red). Right, quantifications. For both **A** and **B**, sample size (*n*) is indicated inside the bars in the graph, each dot represents an embryo. *N* = 4. Phospho-histone H3 (Ser10) intensity was normalized to cytoplasmic levels. The P-values were determined using two-tailed unpaired Student’s t-test. ***p < 0.001, ****p < 0.0001. The dashed lines indicate the outline of embryos. RNA interference was performed by feeding. Scale bar is 10 µm.

Similar to the observation in SDS-22 depleted embryos (Fig 5A), phospho-histone H3 (Ser10) levels were higher in the *sds-22(E153A)* mutant embryos (Fig 5B), suggesting that the interaction between SDS-22 and GSP-1 and GSP-2 is important to maintain efficient dephosphorylation of a PP1 substrate. To conclude, while our genetic data on PAR-2 cortical localization suggest that SDS-22 is not required to fully activate GSP-1 and/or GSP-2, depletion or mutation of SDS-22 results in a reduced activity of the phosphatases, as shown by phospho-histone H3 (Ser10) levels. This suggests that SDS-22 plays a general role in regulating GSP-1 and GSP-2, which is not specific to cell polarity.

### SDS-22 maintains GSP-1 and GSP-2 protein levels by protecting them from proteasomal degradation

Regulatory subunits of phosphatases can regulate their activity, localization or protein levels (Aggen *et al*., 2000; Verbinnen *et al*., 2017; Virshup & Shenolikar, 2009). Recent work in human cells and in vitro has shown that SDS22 coordinates the assembly of the PP1 holoenzyme (Cao *et al*., 2024; Kueck *et al*., 2024). To understand how SDS-22 contributes to the function of GSP-1 and GSP-2, we investigated whether SDS-22 regulated the localization or levels of GSP-1 and GSP-2. We depleted SDS-22 in the *mNG::gsp-2* and *gfp::gsp-1* strains and observed that the localization of GSP-2 and GSP-1 was not altered by SDS-22 depletion (Fig 6A). However, the intensity of mNG::GSP-2 and GFP::GSP-1 dropped to 53.7% and 59.5% compared to the control strains (Fig 6B,C). This result was confirmed by western blot analysis (Fig 6D,E). SDS-22 is therefore essential to maintain the proper levels of GSP-1 and GSP-2 in *C. elegans* embryos.

**Fig 6:**
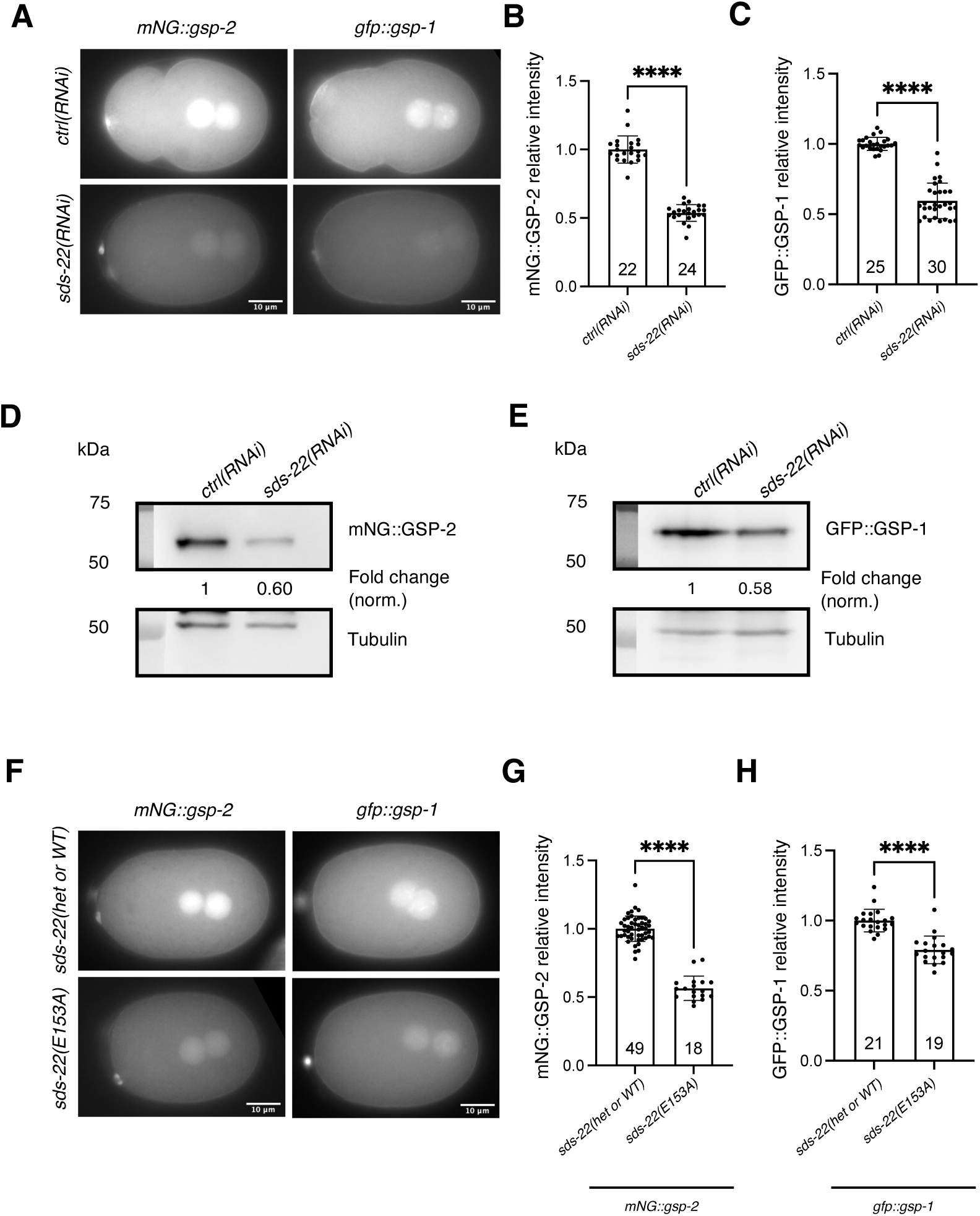
SDS-22 depletion or mutation causes a reduction of GSP-1 and GSP-2 protein levels. **(A)** Representative images of *mNG::gsp-2* and *gfp::gsp-1* embryos in *ctrl(RNAi)* and *sds-22(RNAi)*. **(B, C)** Quantification of relative mNG::GSP-2 **(B)** and GFP::GSP-1 **(C)** levels in *sds-22(RNAi)* normalized to *ctrl(RNAi)*. Mean is shown and error bars indicate SD. *N* = 4. ****p < 0.0001. **(D, E)** Western Blot of embryonic extracts from *mNG::gsp-2* **(D)** and *gfp::gsp-1* **(E)** in *ctrl(RNAi)* and *sds-22(RNAi)* embryos. Tubulin is used as a loading control. Fold change of mNG::GSP-2 and GFP::GSP-1 normalized to Tubulin levels were shown below the Western blot image. *N* = 1. **(F)** Representative images of *mNG::gsp-2; sds-22(E153A/+ or +/+,* a mixture of wild type and heterozygous of the E153A mutant, see methods), *mNG::gsp-2; sds-22(E153A), gfp::gsp-1* and *gfp::gsp-1; sds-22(E153A).* **(G, H)** Quantification of relative mNG::GSP-2 **(G)** and GFP::GSP-1 **(H)** levels with SDS-22(E153A) mutation normalized to wild type. *mNG::gsp-2, N* = 3. *gfp::gsp-1, N* = 4. ****p < 0.0001. In A-E, RNA interference was performed by feeding. In **B**, **C**, **G**, **H**, mean is shown, error bars indicate SD, the P-values were determined using two-tailed unpaired Student’s t-test. Dots in graphs represent individual embryo measurements and sample size (*n*) is indicated inside the bars in the graph.

We next asked whether SDS-22 maintains GSP-1 and GSP-2 levels in vivo via their physical interaction with the proteins. We generated strains in which the E153 site of SDS-22 is mutated in the genetic background of *mNG::gsp-2* and *gfp::gsp-1* separately. In contrast to the low lethality observed in *sds-22(E153A)* mutant in the *gfp::par-2* genetic background (Fig 3C), *sds-22(E153A)* mutation caused 99.1% embryonic lethality in *mNG::gsp-2* and 32.3% in *gfp::gsp-1* (Fig EV6A,B). These data suggest that tagged GSP-2 and GSP-1 strains are hypomorph even though there are no observable growth defects nor embryonic lethality in these strains. Because of the high embryonic lethality of *mNG::gsp-2; sds-22(E153A)* homozygous mutant, we maintained the strain as a *sds-22(E153A)* heterozygote (see methods). Consistent with the phenotypes of *sds-22(RNAi)*, SDS-22(E153A) mutation led to a 46.3% of reduction of the mNG::GSP-2 intensity (Fig 6F,G) and to a 21% reduction of the GFP::GSP-1 intensity (Fig 6F,H).

We then investigated whether the levels of GSP-1 and/or GSP-2 affected the levels of SDS-22. We performed single depletion and co-depletion of GSP-1 and GSP-2 in *sds-22::gfp* strain. We found that the levels of SDS-22 were weakly reduced by single depletion of GSP-1 or GSP-2 (around 8% of reduction in both conditions), but were un-affected by co-depletion of both GSP-1 and GSP-2 (Fig EV7A,B).

In budding yeast, when Sds22 is mutated, PP1-Glc7 is prone to misfold and form aggregates with heat-shock proteins, which are cleared by the proteasome (Cheng & Chen, 2015). Additionally, in human cells, SDS22 stabilizes nascent PP1 (Cao *et al*., 2024). We hypothesized that GSP-1 and GSP-2 are degraded by the proteasome when SDS-22 is depleted or mutated. To test this hypothesis, we reduced proteasomal activity in SDS-22-depleted or mutated embryos by depleting the proteasomal subunits RPN-6.1 or RPN-7, which localize to the 19S regulatory complex and are essential for the proteolytic activity (Fernando *et al*, 2022; Papaevgeniou & Chondrogianni, 2014). Since depletion of these subunits results in worms with very little to no progeny (Fernando *et al*., 2022), we analyzed the protein intensity of GSP-1 and GSP-2 in the –1 and –2 oocytes in the germline and investigated whether reducing proteasomal activity would restore GSP-1 and GSP-2 levels.

When RPN-6.1 or RPN-7 were depleted, there was an increase of GSP-1 and GSP-2 intensity both in the cytoplasm and in the nucleus of the oocytes (Fig 7A-F, Fig EV8), indicating that GSP-1 and GSP-2 are subject to proteasomal degradation in the germline of *C. elegans*. Consistent with the observation in embryos (Fig 6A), both cytoplasmic and nuclear GSP-2 levels in the germline were reduced by about 50% after depletion of SDS-22 (Fig 7B,C). Importantly, when SDS-22 was depleted together with RPN-6.1 or RPN-7, the levels of GSP-2 were restored (Fig 7A-C, Fig EV8A). The efficiency of SDS-22 depletion in the RPN-6.1 or RPN-7 co-depletion was verified (Fig EV9). In *sds-22(RNAi)* oocytes, the levels of GFP::GSP-1 exhibited a smaller decrease (17.4%) in the nucleus and an even smaller reduction in the cytoplasm, which was not statistically significant (8.8%, n.s.) (Fig 7D-F). The reduced nuclear and cytoplasmic intensities of GSP-1 following SDS-22 depletion were also rescued by co-depletion of RPN-6.1 or RPN-7 (Fig 7D-F, Fig EV8B), similar to the recovery of GSP-2 levels (Fig 7B,C). To further test the role of SDS-22 on GSP-1 stability, we measured GFP::GSP-1 levels in the – 1 and –2 oocytes in the *gfp::gsp-1; sds-22(E153A)* strain. In the germline of *gfp::gsp-1; sds-22(E153A)* worms, GSP-1 levels were reduced (28.8% in the cytoplasm; 31.0% in the nucleus) (Fig 7G-I). In the same mutant, depletion of RPN-6.1 or RPN-7 increased the levels of GSP-1 (Fig 7G-I, Fig EV8C), supporting our hypothesis that SDS-22, through its interaction with GSP-1, protects GSP-1 from proteasome mediated degradation. Since we use the embryonic lethality phenotype of the *mNG::gsp-2; sds-22(E153A)* strain to recognize the homozygote *sds-22(E153A),* this precluded the possibility to analyze the germlines of homozygote *mNG::gsp-2; sds-22(E153A)* worms depleted of RNP-6.1 or RPN-7, as these worms do not have progeny (Fernando *et al*., 2022) and we therefore cannot distinguish the *sds-22(E153A)* homozygote from the *sds-22(E153A)* heterozygote (see material and methods for details).

**Fig 7:**
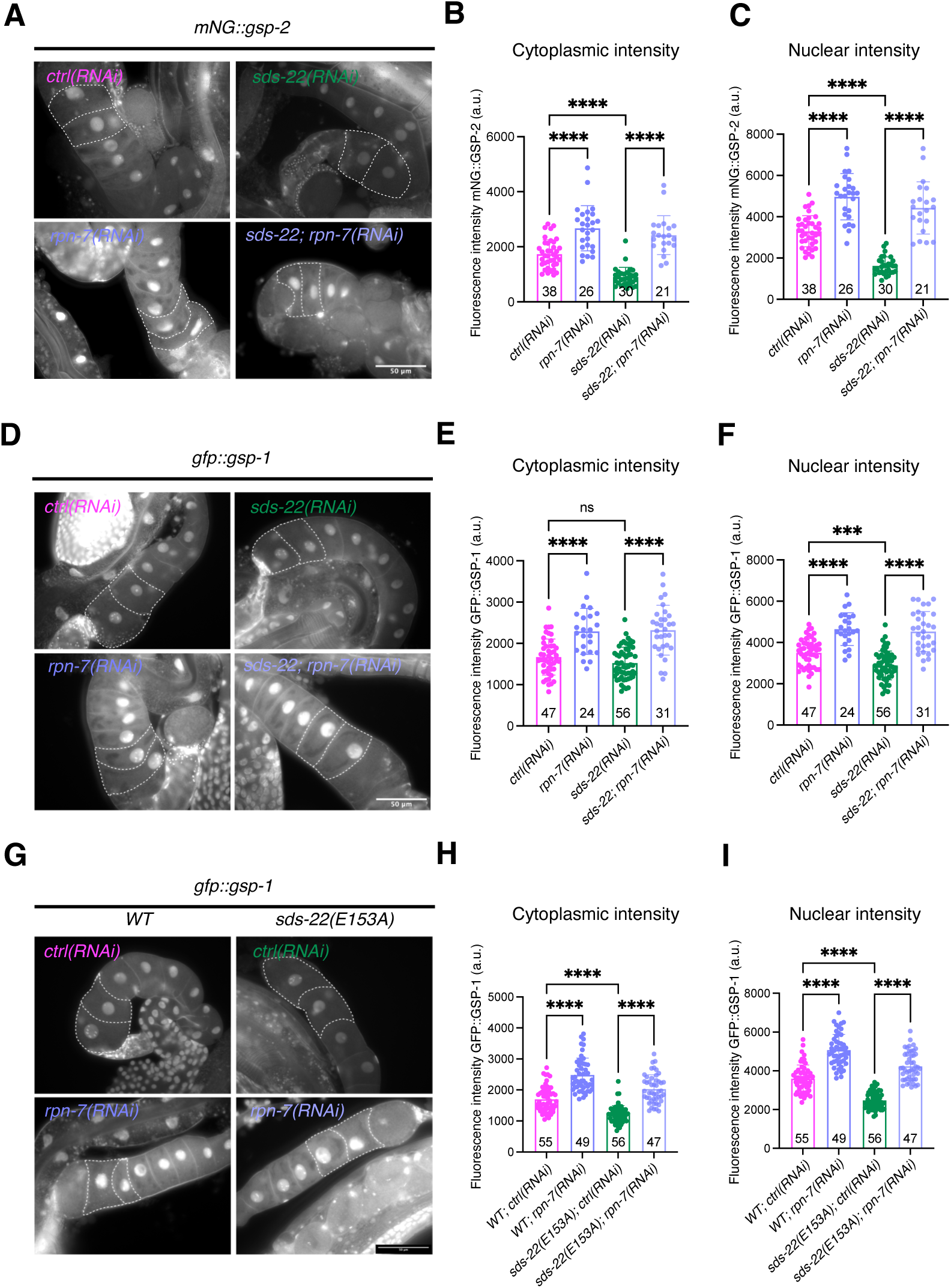
SDS-22 protects GSP-1 and GSP-2 from proteasomal degradation. **(A)** Representative images of *mNG::gsp-2* germlines in *ctrl(RNAi), sds-22(RNAi), rpn-7(RNAi)* and *sds-22(RNAi); rpn-7(RNAi).* **(B, C)** Quantification of mNG::GSP-2 intensity levels in the cytoplasm **(B)** and nucleus **(C)** of –1 and –2 oocytes. *N* = 3. **(D)** Representative images of *gfp::gsp-1* germlines in *ctrl(RNAi), sds-22(RNAi), rpn-7(RNAi)* and *sds-22(RNAi); rpn-7(RNAi).* **(E, F)** Quantification of GFP::GSP-1 intensity levels in the cytoplasm **(E)** and nucleus **(F)** of –1 and –2 oocytes. *N* = 3. **(G)** Representative midsection images of *gfp::gsp-1, gfp::gsp-1;sds-22(E153A)* germlines in *ctrl(RNAi)* and *rpn-7(RNAi).* **(H, I)** Quantification of GFP::GSP-1 intensity levels in the cytoplasm **(H)** and nucleus **(I)** of –1 and –2 oocytes. *N* = 3. For **B**, **C**, **E**, **F**, **H** and **I**, mean is shown and error bars indicate SD. Each dot represents the measurement of one germline. The P-values were determined using one-way ANOVA “Tukey’s multiple comparisons test”. Sample size (*n*) is indicated inside the bars in the graph. For all panels, RNA interference was performed by feeding. The scale bars are 50 µm. In all plots, ns p > 0.05, * p < 0.05, ** p < 0.01, ***p < 0.001, ****p < 0.0001, *n* = number of embryos analyzed; *N* = number of independent experiments.

In summary, these data show that SDS-22 is important to maintain the levels of GSP-1 and GSP-2 by protecting them from proteasome mediated degradation.

## Discussion

Cell polarity establishment in the *C. elegans* embryos is crucial for subsequent asymmetric cell divisions and later development. Previous studies have shown that the balanced activity of the kinase PKC-3 and the PP1 phosphatases GSP-1 and GSP-2 is essential for the posterior cortical localization of PAR-2 (Calvi *et al*., 2022; Hao *et al*., 2006; Motegi *et al*., 2011). GSP-1/-2 are PP1 catalytic subunits that rely on regulatory subunits to direct their cellular localization or facilitate their activity within a functional holoenzyme. Here we find that a conserved PP1 regulator, SDS-22, when depleted, results in a smaller PAR-2 domain and can partially rescue the polarity defects of a *pkc-3(ne4246)* mutant. We demonstrate that SDS-22 contributes to the activity of GSP-1/-2 by protecting them from proteasomal degradation and maintaining their protein levels. Taken together, our data suggest that the role of SDS-22 in polarity is indirect via the regulation of GSP-1/-2 levels. In support of this, SDS-22 depletion results in broader GSP-1/-2 dependent phenotypes such as increased Phospho-H3 (Ser10) (Fig 5) and centriole duplication defects in later-stage embryos (Peel *et al*., 2017).

SDS-22 is a conserved protein previously shown to be important for cell polarity. In *Drosophila,* Sds22 is essential for maintaining the morphology and apical-basal polarity of follicle epithelial cells (Grusche *et al*, 2009) and it facilitates the cortical localization of the lethal giant larvae (Lgl) protein in these cells (Moreira *et al*., 2019). In mammalian and yeast cells, SDS22 is required for mitotic progression (Duan *et al*., 2016; Posch *et al*., 2010; Wurzenberger *et al*., 2012). Even though SDS22 appears as a conserved regulator of PP1 phosphatases, the exact mechanism by which SDS22 regulates PP1 activity was still debated. Some studies have shown that SDS22 positively contributes to PP1 activity in mitosis (MacKelvie *et al*, 1995; Stone *et al*, 1993) and plays a role in localizing PP1 and in protein folding of the PP1 complex (Cheng & Chen, 2015; Peggie *et al*., 2002). However, other studies suggested that SDS22 inhibits PP1 activity in vitro (Lesage *et al*., 2007). Two recent studies from the Bollen and Meyer groups show that SDS22 plays a key role in the biogenesis of PP1 holoenzymes (Cao *et al*., 2024; Kueck *et al*., 2024). On the one hand, SDS22 stabilizes newly translated PP1; on the other hand, SDS22 and Inhibitor 3 (I3) bind to PP1, forming a ternary complex that inhibits PP1 activity. This inhibitory complex is dissociated by the AAA ATPase p97/Valosin, allowing PP1 to bind canonical regulatory subunits to form a functional holoenzyme. Given that SDS-22 both stabilizes PP1 levels and inhibits its activity, this dual role clarifies the apparent contradiction: while SDS-22 is essential for PP1 activity in vivo (because it is essential for the biogenesis/stability), it inhibits PP1 activity in vitro (as it needs to be removed to have an active PP1), while in vivo it is removed by p97/Valosin resulting in active PP1 (Cao *et al*., 2024; Kueck *et al*., 2024).

Our data in *C. elegans* support and complement these studies. We show that SDS-22 is not essential for full activation of both GSP-1 and GSP-2, since depletion of SDS-22 does not abolish polarity, as observed in the co-depletion of GSP-1 and GSP-2 (Calvi *et al*., 2022). Embryos co-depleted of SDS-22 and GSP-1 or of SDS-22 and GSP-2, are also able to polarize, indicating that SDS-22 plays a different role than simply activating GSP-1, GSP-2 or both. We find that in *C. elegans* SDS-22 is required to maintain the appropriate levels of both GSP-1 and GSP-2. We propose that the reduced levels of GSP-2 explain why loss of SDS-22 partially rescues PAR-2 defect in *pkc-3(ne4246)* mutant allele on one side (Fig 1B). On the other side, since embryos still have GSP-1 and GSP-2 proteins they are able to polarize after the depletion of SDS-22, or co-depletion of SDS-22 with GSP-1 or SDS-22 with GSP-2 in *gfp::par-2* embryos (Fig 4). In these conditions, depletion of SDS-22, whether alone or in conjunction with GSP-1 or GSP-2, only results in a partial loss of the PP1 subunits GSP-1 and/or GSP-2 and in a reduction in catalytic activity. Consistent with GSP-2 reduced levels, SDS-22 depleted or E153A mutant embryos also have a smaller PAR-2 domain. However, since these embryos also show reduced cortical ruffling (Movie EV1,2) and are smaller (Fig EV2C) we cannot exclude that these two phenotypes also contribute to the smaller size of the PAR-2 domain.

How SDS-22 maintains PP1 level has not been fully clarified. In budding yeast, it has been reported that when Sds22 is mutated, PP1/Glc7 forms aggregates in the nucleus, despite the overall protein level remaining unaltered (Peggie *et al*., 2002). These aggregates co-localize with heat-shock proteins that require clearance by the proteasome (Cheng & Chen, 2015). Co-expression of PP1 with chaperones GroEL/ES(HSP-60) in bacteria (Peti *et al*, 2013) or with SDS22 in mammalian cells (Choy *et al*, 2019) increases PP1 yield and solubility. These findings suggest that in the absence of SDS22, PP1 is prone to mislocalization or misfolding, leading to aggregation. Interestingly, Peel et al. found SDS-22 co-immunoprecipitated with TCP-1 and HSP-60 (Peel *et al*., 2017), both members of the chaperonin family critical for protein folding (Hayer-Hartl *et al*, 2016; Kubota *et al*, 1995). Collectively, these findings suggest that by facilitating protein folding SDS22 protects PP1 from degradation. In the *C. elegans* germline and embryos GSP-1/-2 do not form visible aggregates when SDS-22 is depleted but they undergo degradation in a proteasome-dependent manner (Fig 7). Our data, together with previous findings (Cao *et al*., 2024; Cheng & Chen, 2015), demonstrate a role of SDS-22 in protecting PP1s from proteasomal degradation. Interestingly, E153A mutation of SDS-22 affected GSP-2 levels much more strongly than GSP-1 levels in embryos (Fig 6F-H). Furthermore, in germlines, GSP-1 levels were less affected by loss of SDS-22 (Fig 7). This indicates that GSP-2 is more susceptible to the interaction with SDS-22.

Our data and the recent studies of the Bollen and Meyer laboratories (Cao *et al*., 2024; Kueck *et al*., 2024) suggest that there is a canonical regulator of GSP-1 and GSP-2 in embryonic polarity that we have not yet identified. In the immunoprecipitations of GSP-2 from embryos we have identified APE-1 and SZY-2(I-2). APE-1 has been shown to direct GSP-2, but not GSP-1, localization to epidermal junctions and modulate GSP-2 activity (Beacham *et al*., 2022). SZY-2(I-2) regulates both GSP-1/-2 in centriole duplication from the four-cell stage of *C. elegans* embryos (Peel *et al*., 2017). However, our data suggest that neither APE-1 nor SZY-2 functions as canonical PP1 regulators in PAR-2’s regulation, as depletion of either did not rescue the polarity defects of *pkc-3(ne4246)* mutant (Fig 1A, EV2A). We cannot exclude the possibility that SDS-22 may also function as a canonical regulator of the PP1 holoenzymes in controlling PAR-2 localization in *C. elegans* embryos. Further investigation is required to determine whether a yet-to-be-identified regulator of the PP1 holoenzyme is responsible for loading PAR-2 to the posterior cortex.

Another open question is if and how the activity of PP1 catalytic subunits is asymmetrically modulated in the one-cell embryo. Our previous work (Calvi *et al*., 2022) suggests that the activity of GSP-1/-2 might be asymmetrically restricted by the anterior-enriched polo-like kinase PLK-1, which consequently gives rise to asymmetric posterior localization of PAR-2. One possibility is that PLK-1 directly phosphorylates and regulates the activity of GSP-1 and/or GSP-2. Interestingly, a recent study shows that in *Drosophila* cells POLO kinase phosphorylates and inhibits PP1 (Moura *et al*, 2024). However, we were not able as yet to identify potential phosphorylated sites that regulates PP1 activity (our unpublished data). Another possibility is that PLK-1 regulates the regulatory subunit(s) of GSP-1/-2. Notably, in mammalian cells, SDS22 is phosphorylated and inhibited by Plk1 during mitosis (Duan *et al*., 2016). Further studies will be required to understand whether additional pathways regulate the activity of this phosphatase during polarity establishment.

## Material and Methods

### Worm strains

Worms were maintained on NGM plates seeded with OP50 bacteria using standard methods (Brenner, 1974). The strains used in this work are listed in Table S1. The strains containing the temperature sensitive *pkc-3(ne4246)* mutation, *pkc-3(ne4246); gfp::par-2* (Ng *et al*, 2022) and *pkc-3(ne4246); gfp::par-2; sds-22(E153A),* were kept at 15°C. All the other strains were maintained at 20°C.

Mutant strains were generated using CRISPR/Cas9 technology as described (Arribere *et al*, 2014). The single guide RNAs, repair templates used to generate the mutants, PCR primers used to amplify the sequence with mutation, as well as enzymes used for the screening, are listed in Table S2-4. For the strain ZU341 *(gfp::par-2; sds-22(E153A))*, two independent isolates were analyzed. The ZU362 *mNG::gsp-2; sds-22(E153A)* was maintained as a *sds-22(E153A)* heterozygote. *mNG::gsp-2; sds-22(E153A)* adult worms are viable but produce 100% dead progeny, indicating that this mutation behaves as a maternal effect lethal in the *mNG::gsp-2* background.

### RNA interference

Clones from the Ahringer feeding library (Ahringer, 2006; Kamath *et al*, 2003) were used as listed in Table S5. The clone C06A6.2 served as the control for the injection experiments, as previously done (Bondaz *et al*, 2019). A fragment of APE-1 was amplified from cDNA and cloned into the final pDEST-L4440 vector using Gateway technology (primer listed in Table S5). Double-stranded RNA (dsRNA) for injections was produced with the Promega Ribomax RNA production system. dsRNA was injected in L4/young adult hermaphrodites, which were then incubated at 20°C. For the co-depletion of SDS-22 and GSP-1/-2, and co-depletion of GSP-1 and GSP-2, a mixture of 1:1 dsRNAs was injected (Fig 4, Fig EV5, EV7). Embryos from injected worms were analyzed 16-20 hours after injection.

RNA interference by feeding was performed on plates containing 1mM IPTG. The L4440 vector was used as control. For the *pkc-3(ne4246)* strains and controls, worms were incubated on feeding plates at semi-restrictive temperatures of 24°C (Fig 1, Fig 3D, Fig EV2A) or 22°C (Fig 3E,F,G). L4 larvae were added to RNAi feeding plates and incubated for 24 hours. For co-depletion of SDS-22 and RPN-6.1/-7 (Fig 7, EV8-9), a mixture of 1:1 fresh bacteria culture was plated. Worms were incubated on feeding plates at 20°C for 24h hours (Fig 5A, Fig 6A-E, Fig7, EV2B, EV3, EV8-9).

### Immunoprecipitation and mass spectrometry

For immunoprecipitation experiments from embryos of mNeonGreen tagged GSP-2, worms were grown in 3X PEP plates and embryos were harvested from gravid worms by bleaching (500 mM NaOH and 5% bleach). The *N2* strain was used as a control. The mNeonGreen::GSP-2 and control immunoprecipitations were performed in triplicate. Packed embryos were frozen in liquid nitrogen and stored at −80°C. Embryos were then thawed, resuspended in immunoprecipitation buffer (100 mM KCl, 50 mM Tris (pH: 7.5), 1 mM MgCl2, 1 mM DTT, 5% glycerol, 0.2% NP-40, 1 mM EDTA, and protease and phosphatase inhibitor cocktail (Roche)). The embryos were ground using a mill (Retsch MM301) and silica beads (Lysing matrix C, MP Biomedicals). The protein homogenate was centrifuged at 14,000 rpm for 30 min at 4°C. The protein concentration in the supernatant was determined in a Bradford assay (Bio-Rad Laboratories) using a UV/Vis Spectrophotomer (Labgene scientific). About 3 mg of embryonic extract was incubated at 4°C for 1.5 h with 10 µl of mNeonGreen-Trap Agarose beads (Chromotek), previously washed 3 times with IP buffer. After the incubation with the embryonic extract, beads were washed three times with IP buffer, and three times with the buffer containing 100 mM KCl, 50 mM Tris (pH: 7.5), 1 mM MgCl2, 1 mM DTT, 1 mM EDTA to remove nonspecific binding. Elution of mNeonGreen-tagged GSP-2 was performed by incubation of the beads for 10min at room temperature with 25 µl of 0.15% TFA. The pH of the elution was corrected to pH: 7-8 with 0.5M Tris (pH:7.9). 3 µl of the elutions were used to analyse the immunoprecipitation by western blot. 22 µl of the elutions were snap frozen and sent to Biognosys (https://biognosys.com) for mass spectrometric analysis. The elutions were subjected to denaturation, reduction, alkylation, digestion and C18 clean up. The peptide digests were acquired in DIA mode using an Orbitrap Exploris 480 (Thermo Fisher) and analyzed with directDIA (Spectronaut 15).

### Imaging of live embryos

Adult hermaphrodites were dissected on a coverslip in a drop of Egg Buffer (118 mM NaCl, 48 mM KCl, 2 mM CaCl₂, 2 mM MgCl₂, and 25 mM Hepes, pH 7.5). Embryos were mounted on a 3% agarose pad for imaging. Time-lapse recordings, with frames captured every 10 seconds, were conducted using a Nikon ECLIPSE Ni-U microscope equipped with a Nikon DS-U3 Digital Camera and a 60X/1.25 numerical aperture (NA) objective. Imaging was performed between 20 – 22°C.

### Immunostaining and imaging of fixed embryos

Fixation and staining of embryos were carried out as described previously (Calvi *et al*., 2022). The primary antibody phospho-Histone 3 (Ser10) (Millipore, rabbit) was diluted 1:500 in PBST with 1% BSA and slides incubated overnight at 4°C. Slides were then washed and incubated with the secondary antibody (4 μg/ml Alexa Fluor 568–coupled anti-rabbit antibody) and 1 μg/ml DAPI to visualize DNA. Images were acquired using a Nikon ECLIPSE Ni-U microscope, equipped with a Nikon DS-U3 Digital Camera, and using a 60X/1.25 numerical aperture (NA) objective.

### Embryonic lethality

To assess the embryonic lethality, young adult worms were singled on NGM plates seeded with OP50 and incubated at 20°C for 24 hours. Adults were the removed and the plates were incubated for an additional 24 hours at 20°C (Fig 3C, Fig EV6, and other *sds-22(E153A)* mutants in *N2* and in *sds-22::gfp* genetic backgrounds). The *mNG::gsp-2; sds-22(E153A)* strain was maintained as a *sds-22(E153A)* heterozygote, since in the homozygote form it is 99.1% embryonic lethal. To quantify the lethality of the homozygote, L4/young adult worms (homozygote wild-type, heterozygote, or homozygote *sds-22(E153A)* mutant) were singled on OP50 seeded-NGM plates, allowed to lay eggs for 24 hours, and subsequently lysed for genotyping. Homozygote *sds-22* wild-type and heterozygote worms expressing wildtype *sds-22* from one allele and mutant *sds-22(E153A)* from the other allele were used as a control. The embryonic lethality can be used to differentiate homozygotes from heterozygotes and wild types for quantification (e.g. Fig 6F,G). However, since depletion of RNP-6.1 or RPN-7 resulted in few or no progeny (Fernando *et al*., 2022), this precluded the possibility to analyze the germlines of homozygote *mNG::gsp-2; sds-22(E153A)* worms with RNP-6.1 or RPN-7 depletion, as we cannot distinguish the homozygote from the heterozygote.

SDS-22(E153A) mutation in wildtype *N2* exhibited 8.3% of lethality, while *sds-22(E153A)::gfp* showed a lethality of 51.9%.

For the RNAi feeding and lethality assay of *pkc-3(ne4246)* mutant, L4 worms were initially transferred to 1mM IPTG plates seeded with feeding bacteria and incubated overnight at 15°C. The following day, young adult worms were singled onto feeding plate and incubated for 24 hours at 24°C (Fig 1A, 3D) and 22°C (Fig 3E). After this period, the adult worms were removed and the plates were further incubated at 24°C and 22°C respectively for 24 hours. The ratio of unhatched embryos to the total F1 progeny (unhatched embryos plus larvae) was used to calculate the percentage of embryonic lethality.

### Yeast two-hybrid assay

The interaction between SDS-22 and GSP-1/-2 was assessed using a GAL4-based yeast two-hybrid (Y2H) system (Gateway, Invitrogen) using the MAV203 yeast strain. Full-length SDS-22 (1-326) fragments, both wild type and mutant (E153A), were fused to the GAL4 DNA binding domain (DBD, Bait plasmid). Full-length cDNAs of GSP-1 and GSP-2 were fused to the GAL4 activation domain (AD, Prey plasmid). The SDS-22 wild type and mutant fragments, GSP-1 and GSP-2 were first cloned into the pDONR201, then transferred to the pDEST32 (GAL4DBD) and pDEST22 vector (GAL4AD) separately using Gateway technology. Mutations were introduced by Pfu site-directed mutagenesis. A list of plasmids and primers used for the Y2H assay is provided in Table S6 and S7 respectively. Transformants were selected on synthetic defined (SD) medium lacking leucine and tryptophan. Interactions were tested by spotting single colonies containing the desired plasmids on medium lacking leucine, tryptophan and histidine, with 7.5mM of 3AT (3-amino-1,2,3-triazole, Sigma). Picture of the plates were captured using the Fusion FX6 EDGE Imaging System (Vilber) equipped with an Evo-6 Scientific Grade CCD camera.

### Western blot

Embryos obtained by hypochlorite treatment from two 5cm plates of adult worms were resuspended in SDS sample buffer and denatured at 95°C for 5 minutes. Approximately 3000 embryos per sample were loaded onto a 10% SDS-PAGE gel, followed by Western blotting with ECL detection (Vilber, Fusion FX). Primary antibodies [(anti-TUBULIN (1/2500, mouse, Sigma), anti-mNeonGreen (1/1000, mouse, ChromoTek) and anti-GFP (1/2500, rabbit, Pines)] were incubated overnight at 4°C. HRP-conjugated secondary antibodies were incubated for 45 minutes at room temperature.

Band signal measurement for anti-TUBULIN, anti-mNeonGreen, and anti-GFP was performed in Fiji ImageJ by drawing equal-sized ROIs over each band. Mean intensity values were used for quantification, and background was subtracted using the mean intensity of an equivalent ROI placed below the band of interest. The ratio of *ctrl(RNAi)* treatment was normalized to 1.

### Image analysis and measurement

#### Cortical PAR-2 intensity measurement (Fig1 B,C, Fig 3F,G)

The line file of GFP::PAR-2 intensity along the entire cortex was measured in one-cell stage embryos at pronuclear meeting. The stage of pronuclear meeting is defined when the two pronuclei first contact each other. In *pkc-3(ne4246)* embryos, the two pronuclei exhibited a tendency to meet more centrally compared to controls (Fig 1B, Movie EV1), as shown in (Kirby *et al*, 1990; Rodriguez *et al*, 2017). Using ImageJ software, a free-line 5-pixel-wide was positioned at the anterior pole of the cortex (position 0 μm) and traced manually along the whole cortex to the posterior pole (position 75 μm) and back to the anterior pole. The background was subtracted from each value. The intensity of the line profile (from 0 to 75 μm) of the segment traced was shown in an arbitrary unit (a.u.). Images were in 16-bit format.

#### Cortical PAR-2 size measurement (Fig 1D, Fig 3B, Fig 4)

The size of cortical GFP::PAR-2 domain was determined by measuring the length of the PAR-2 domain normalized for the total perimeter of the embryo at pronuclear meeting. The perimeter of the embryo and the length of the PAR-2 domain were measured by the Qupath software with machine learning, as detailed previously (Vaudano *et al*, 2024). The length of the PAR-2 domain is represented as a percentage of the total perimeter of the embryo.

#### AB cell size measurement (Fig 1E,F)

The total length of the embryo was measured manually from the anterior to the posterior pole using ImageJ software. AB cell size was measured from the anterior pole to the cleavage midline when the embryo just divided into two cells. The AB cell ratio was calculated as the proportion of AB cell size relative to the total embryo length (AB cell plus P1 cell length).

#### Protein intensity of mNG::GSP-2, GFP::GSP-1, SDS-22::GFP in embryos and germlines (*Fig 6-7*, Fig EV4, Fig EV7-9)

For embryos, the mean intensity of cytoplasmic mNG::GSP-2, GFP::GSP-1, SDS-22::GFP was determined by tracing a fixed-length line in the cytoplasm from the anterior to the posterior of the embryo (above the nucleus) at pronuclear meeting stage using the ImageJ software. The background was subtracted from each value.

For the embryonic intensity of mNG::GSP-2 in the *sds-22(E153A)* mutant genetic background, which is embryonic lethal, L4/young adult worms (homozygote wild type, heterozygote or homozygote *mNG::GSP-2; sds-22(E153A)* mutant) were singled, let them lay few eggs, and subsequently dissected for imaging. Homozygous *mNG::GSP-2; sds-22(E153A)* mutant worms were recognized by looking at the progeny in the original plates, as homozygote *mNG::GSP-2; sds-22(E153A)* worms only lay dead eggs. Homozygote *sds-22* wild-type and heterozygote worms expressing wildtype *sds-22* from one allele and mutant *sds-22(E153A)* from the other allele were used as a control.

For germlines, mean cytoplasmic and nuclear intensities of mNG::GSP-2, GFP::GSP-1, SDS-22::GFP were measured by drawing a fixed circle in the cytoplasm and/-or nucleus of –1 and –2 oocytes using ImageJ. Background intensity was subtracted. The mean intensities of the cytoplasmic and nuclear regions in –1 and –2 oocytes were calculated and reported.

The germline intensity of mNG::GSP-2 in the *sds-22(E153A)* mutant cannot be measured because there were no viable embryos after depletion of RNP-6.1 or RNP-7, therefore the genotype of homozygous *mNG::gsp-2; sds-22(E153A)* (also embryonic lethal) cannot be differentiated from the homozygote wild type and heterozygote.

#### Phospho-Histone 3 (Ser10) intensity measurement (*Fig 5*)

One-cell stage embryos at anaphase were analyzed. A segment 5-pixel-wide and of constant length, centered at the chromosomes was traced using ImageJ software. For each embryo, the average of three segments (upper, center, and lower chromosomes) was used for quantification. The intensity of the line profile of each embryo was normalized to the average of the value in the cytoplasm at position 0 of the segment traced. The mean intensities of the phospho-Histone 3 (Ser10) on the chromosomes were calculated and reported.

## Statistical analysis

Statistical analysis was conducted using GraphPad Prism 9. Details on the statistical tests, sample sizes, number of replicates, and interpretation of error bars are specified in each figure legend, within the results section, or in the methods section, and summarized in Table S8. Data was assumed to have normal distribution.

Significance was defined as: ns, p > 0.05, *p < 0.05, **p < 0.01, ***p < 0.001, ****p < 0.0001.

## Data availability

All the strains and reagents generated in this study are available from the corresponding author upon request. All raw data associated with the experiments will be deposited in Yareta at publication.

## Author Contributions

Yi Li: Conceptualization; Formal analysis; Investigation; Visualization; Writing-original draft; Writing-review and editing. Ida Calvi: Conceptualization; Investigation; Writing-review and editing. Monica Gotta: Conceptualization; Funding acquisition; Supervision; Writing-original draft; Writing-review and editing

## Acknowledgments

A special thanks to Erik Griffin, and Dhanya Cheerambathur for strains. We would like to thank all the present and past members of the Gotta, Meraldi and Steiner laboratories, Patrick Meraldi and Florian Steiner for discussions and suggestions. Thanks to Sofia Barbieri, Victoria von Glasenapp and Florian Steiner for comments on the manuscript. Some strains were provided by the CGC, which is funded by NIH Office of Research Infrastructure Programs (P40 OD010440). This research was funded by the Swiss National Science Foundation (grant number 31003A_175850) and the University of Geneva.

## Supplementary movies

**Movie EV1: *sds-22(RNAi)* rescues PAR-2 posterior cortical localization in the *pkc-3(ne4246); gfp::par-2* embryos.** *gfp::par-2* and *pkc-3*(*ne4246*); *gfp::par-2* embryos treated either with *ctrl*(*RNAi*) or *sds-22*(*RNAi*) (RNAi by feeding). Acquisition of midplane fluorescent images begins during the early establishment phase, and frames are captured every 10 s. In *gfp::par-2* embryos (both *ctrl*(*RNAi*) and *sds-22*(*RNAi*)) PAR-2 localizes at the posterior cortex in the one-cell stage embryo and in P1 in the two-cell stage embryo (*n* = 12 for both). In *pkc-3*(*ne4246*); *gfp::par-2*, *ctrl*(*RNAi*) PAR-2 is uniformly distributed at the cortex in both one– and two-cell stage embryo (*n* = 42). After depletion of SDS-22, *pkc-3*(*ne4246*); *gfp::par-2* embryos showed PAR-2 posterior localization in one-cell stage embryo and in the P1 cell (*n* = 42 out of 48). *N* = 5. *n* = number of embryos analyzed; *N* = number of independent experiments. Anterior is to the left and posterior to the right. Referred to Fig 1B.

**Movie EV2: *sds-22*(*E153A*) substitution reduces the length of the PAR-2 cortical domain.** Acquisition of midplane fluorescent images begins during the early stage, and frames are captured every 10 s. In *gfp::par-2; sds-22(E153A)* mutant embryos the length of cortical PAR-2 domain is decreased compared to *gfp::par-2* embryos at pronuclear meeting stage (*n* = 30 and *n* = 52, respectively). *N* = 3. *n* = number of embryos analyzed. *N* = number of independent experiments. Anterior is to the left and posterior to the right. Referred to Fig 3A,B.

**Movie EV3: *sds-22*(*E153A*) substitution rescues PAR-2 posterior cortical localization in the *pkc-3*(*ne4246*); *gfp::par-2* embryos at 22°C.** Acquisition of midplane fluorescent images begins during the early stage, and frames are captured every 10 s. In *gfp::par-2; pkc-3*(*ne4246*); *sds-22(E153A)* mutant embryos PAR-2 is more enriched to the posterior cortex compared to *gfp::par-2; pkc-3*(*ne4246*) embryos where PAR-2 is localized all over the cortex (*n* = 19 for both genotypes). *N* = 3. *n* = number of embryos analyzed. *N* = number of independent experiments. Anterior is to the left and posterior to the right. Referred to Fig 3F,G.

## References

1. Aggen JB, Nairn AC, Chamberlin R (2000) Regulation of protein phosphatase-1. Chem Biol 7: R13–23

2. Ahringer J (2006) Reverse Genetics. WormBook: The Online Review of C. elegans Biology [Internet]. WormBook; 2005-2018.

3. Arribere JA, Bell RT, Fu BX, Artiles KL, Hartman PS, Fire AZ (2014) Efficient marker-free recovery of custom genetic modifications with CRISPR/Cas9 in Caenorhabditis elegans. Genetics 198: 837–846

4. Beacham GM, Wei DT, Beyrent E, Zhang Y, Zheng J, Camacho MMK, Florens L, Hollopeter G (2022) The Caenorhabditis elegans ASPP homolog APE-1 is a junctional protein phosphatase 1 modulator. Genetics 222

5. Billmyre KK, Doebley AL, Spichal M, Heestand B, Belicard T, Sato-Carlton A, Flibotte S, Simon M, Gnazzo M, Skop A et al (2019) The meiotic phosphatase GSP-2/PP1 promotes germline immortality and small RNA-mediated genome silencing. PLoS Genet 15: e1008004

6. Bondaz A, Cirillo L, Meraldi P, Gotta M (2019) Cell polarity-dependent centrosome separation in the C. elegans embryo. J Cell Biol 218: 4112–4126

7. Brenner S (1974) The genetics of Caenorhabditis elegans. Genetics 77: 71–94

8. Calvi I, Schwager F, Gotta M (2022) PP1 phosphatases control PAR-2 localization and polarity establishment in C. elegans embryos. J Cell Biol 221

9. Campanale JP, Sun TY, Montell DJ (2017) Development and dynamics of cell polarity at a glance. J Cell Sci 130: 1201–1207

10. Cao X, Lake M, Van der Hoeven G, Claes Z, Del Pino Garcia J, Lemaire S, Greiner EC, Karamanou S, Van Eynde A, Kettenbach AN et al (2024) SDS22 coordinates the assembly of holoenzymes from nascent protein phosphatase-1. Nat Commun 15: 5359

11. Cao X, Lemaire S, Bollen M (2022) Protein phosphatase 1: life-course regulation by SDS22 and Inhibitor-3. FEBS J 289: 3072–3085

12. Ceulemans H, Bollen M (2004) Functional diversity of protein phosphatase-1, a cellular economizer and reset button. Physiol Rev 84: 1–39

13. Ceulemans H, Vulsteke V, De Maeyer M, Tatchell K, Stalmans W, Bollen M (2002) Binding of the concave surface of the Sds22 superhelix to the alpha 4/alpha 5/alpha 6-triangle of protein phosphatase-1. J Biol Chem 277: 47331–47337

14. Cheng YL, Chen RH (2015) Assembly and quality control of the protein phosphatase 1 holoenzyme involves the Cdc48-Shp1 chaperone. J Cell Sci 128: 1180–1192

15. Choy MS, Moon TM, Ravindran R, Bray JA, Robinson LC, Archuleta TL, Shi W, Peti W, Tatchell K, Page R (2019) SDS22 selectively recognizes and traps metal-deficient inactive PP1. Proc Natl Acad Sci U S A 116: 20472–20481

16. Duan H, Wang C, Wang M, Gao X, Yan M, Akram S, Peng W, Zou H, Wang D, Zhou J et al (2016) Phosphorylation of PP1 Regulator Sds22 by PLK1 Ensures Accurate Chromosome Segregation. J Biol Chem 291: 21123–21136

17. Eiteneuer A, Seiler J, Weith M, Beullens M, Lesage B, Krenn V, Musacchio A, Bollen M, Meyer H (2014) Inhibitor-3 ensures bipolar mitotic spindle attachment by limiting association of SDS22 with kinetochore-bound protein phosphatase-1. EMBO J 33: 2704–2720

18. Fernando LM, Quesada-Candela C, Murray M, Ugoaru C, Yanowitz JL, Allen AK (2022) Proteasomal subunit depletions differentially affect germline integrity in C. elegans. Front Cell Dev Biol 10: 901320

19. Goehring NW (2014) PAR polarity: from complexity to design principles. Exp Cell Res 328: 258–266

20. Goldstein B, Macara IG (2007) The PAR proteins: fundamental players in animal cell polarization. Dev Cell 13: 609–622

21. Grusche FA, Hidalgo C, Fletcher G, Sung HH, Sahai E, Thompson BJ (2009) Sds22, a PP1 phosphatase regulatory subunit, regulates epithelial cell polarity and shape [Sds22 in epithelial morphology]. BMC Dev Biol 9: 14

22. Hao Y, Boyd L, Seydoux G (2006) Stabilization of cell polarity by the C. elegans RING protein PAR-2. Dev Cell 10: 199–208

23. Hayer-Hartl M, Bracher A, Hartl FU (2016) The GroEL-GroES Chaperonin Machine: A Nano-Cage for Protein Folding. Trends Biochem Sci 41: 62–76

24. Heroes E, Van der Hoeven G, Choy MS, Garcia JDP, Ferreira M, Nys M, Derua R, Beullens M, Ulens C, Peti W et al (2019) Structure-Guided Exploration of SDS22 Interactions with Protein Phosphatase PP1 and the Splicing Factor BCLAF1. Structure 27: 507–518 e505

25. Hsu JY, Sun ZW, Li X, Reuben M, Tatchell K, Bishop DK, Grushcow JM, Brame CJ, Caldwell JA, Hunt DF et al (2000) Mitotic phosphorylation of histone H3 is governed by Ipl1/aurora kinase and Glc7/PP1 phosphatase in budding yeast and nematodes. Cell 102: 279–291

26. Kamath RS, Fraser AG, Dong Y, Poulin G, Durbin R, Gotta M, Kanapin A, Le Bot N, Moreno S, Sohrmann M et al (2003) Systematic functional analysis of the Caenorhabditis elegans genome using RNAi. Nature 421: 231–237

27. Kapoor S, Kotak S (2019) Centrosome Aurora A regulates RhoGEF ECT-2 localisation and ensures a single PAR-2 polarity axis in C. elegans embryos. Development 146

28. Kirby C, Kusch M, Kemphues K (1990) Mutations in the par genes of Caenorhabditis elegans affect cytoplasmic reorganization during the first cell cycle. Dev Biol 142: 203–215

29. Klinkert K, Levernier N, Gross P, Gentili C, von Tobel L, Pierron M, Busso C, Herrman S, Grill SW, Kruse K et al (2019) Aurora A depletion reveals centrosome-independent polarization mechanism in Caenorhabditis elegans. Elife 8

30. Kubota H, Hynes G, Willison K (1995) The chaperonin containing t-complex polypeptide 1 (TCP-1). Multisubunit machinery assisting in protein folding and assembly in the eukaryotic cytosol. Eur J Biochem 230: 3–16

31. Kueck AF, van den Boom J, Koska S, Ron D, Meyer H (2024) Alternating binding and p97-mediated dissociation of SDS22 and I3 recycles active PP1 between holophosphatases. Proc Natl Acad Sci U S A 121: e2408787121

32. Lang CF, Munro E (2017) The PAR proteins: from molecular circuits to dynamic self-stabilizing cell polarity. Development 144: 3405–3416

33. Lesage B, Beullens M, Pedelini L, Garcia-Gimeno MA, Waelkens E, Sanz P, Bollen M (2007) A complex of catalytically inactive protein phosphatase-1 sandwiched between Sds22 and inhibitor-3. Biochemistry 46: 8909–8919

34. Llense F, Etienne-Manneville S (2015) Front-to-Rear Polarity in Migrating Cells. In: Cell Polarity 1: Biological Role and Basic Mechanisms, Ebnet K. (ed.) pp. 115–146. Springer International Publishing: Cham

35. Longhini KM, Glotzer M (2022) Aurora A and cortical flows promote polarization and cytokinesis by inducing asymmetric ECT-2 accumulation. Elife 11

36. MacKelvie SH, Andrews PD, Stark MJ (1995) The Saccharomyces cerevisiae gene SDS22 encodes a potential regulator of the mitotic function of yeast type 1 protein phosphatase. Mol Cell Biol 15: 3777–3785

37. Meiselbach H, Sticht H, Enz R (2006) Structural analysis of the protein phosphatase 1 docking motif: molecular description of binding specificities identifies interacting proteins. Chem Biol 13: 49–59

38. Moreira S, Osswald M, Ventura G, Goncalves M, Sunkel CE, Morais-de-Sa E (2019) PP1-Mediated Dephosphorylation of Lgl Controls Apical-basal Polarity. Cell Rep 26: 293–301 e297

39. Motegi F, Zonies S, Hao Y, Cuenca AA, Griffin E, Seydoux G (2011) Microtubules induce self-organization of polarized PAR domains in Caenorhabditis elegans zygotes. Nat Cell Biol 13: 1361–1367

40. Moura M, Barbosa J, Pinto I, Leça N, Cunha-Silva S, Verza AE, Pedroso PD, Lemaire S, Duro J, Silva M et al (2024) POLO kinase inhibits Protein Phosphatase 1 to promote the Spindle Assembly Checkpoint and prevent aneuploidy. bioRxiv: 2024.2012.2023.630043

41. Munro E, Nance J, Priess JR (2004) Cortical flows powered by asymmetrical contraction transport PAR proteins to establish and maintain anterior-posterior polarity in the early C. elegans embryo. Dev Cell 7: 413–424

42. Ng K, Bland T, Hirani N, Goehring NW (2022) An analog sensitive allele permits rapid and reversible chemical inhibition of PKC-3 activity in C. elegans. MicroPubl Biol 2022

43. Papaevgeniou N, Chondrogianni N (2014) The ubiquitin proteasome system in Caenorhabditis elegans and its regulation. Redox Biol 2: 333–347

44. Pedelini L, Marquina M, Arino J, Casamayor A, Sanz L, Bollen M, Sanz P, Garcia-Gimeno MA (2007) YPI1 and SDS22 proteins regulate the nuclear localization and function of yeast type 1 phosphatase Glc7. J Biol Chem 282: 3282–3292

45. Peel N, Iyer J, Naik A, Dougherty MP, Decker M, O’Connell KF (2017) Protein Phosphatase 1 Down Regulates ZYG-1 Levels to Limit Centriole Duplication. PLoS Genet 13: e1006543

46. Peggie MW, MacKelvie SH, Bloecher A, Knatko EV, Tatchell K, Stark MJ (2002) Essential functions of Sds22p in chromosome stability and nuclear localization of PP1. J Cell Sci 115: 195–206

47. Peti W, Nairn AC, Page R (2013) Structural basis for protein phosphatase 1 regulation and specificity. FEBS J 280: 596–611

48. Posch M, Khoudoli GA, Swift S, King EM, Deluca JG, Swedlow JR (2010) Sds22 regulates aurora B activity and microtubule-kinetochore interactions at mitosis. J Cell Biol 191: 61–74

49. Qian J, Lesage B, Beullens M, Van Eynde A, Bollen M (2011) PP1/Repo-man dephosphorylates mitotic histone H3 at T3 and regulates chromosomal aurora B targeting. Curr Biol 21: 766–773

50. Riga A, Castiglioni VG, Boxem M (2020) New insights into apical-basal polarization in epithelia. Curr Opin Cell Biol 62: 1–8

51. Rodrigues NT, Lekomtsev S, Jananji S, Kriston-Vizi J, Hickson GR, Baum B (2015) Kinetochore-localized PP1-Sds22 couples chromosome segregation to polar relaxation. Nature 524: 489–492

52. Rodriguez J, Peglion F, Martin J, Hubatsch L, Reich J, Hirani N, Gubieda AG, Roffey J, Fernandes AR, St Johnston D et al (2017) aPKC Cycles between Functionally Distinct PAR Protein Assemblies to Drive Cell Polarity. Dev Cell 42: 400–415 e409

53. Santoro A, Vlachou T, Carminati M, Pelicci PG, Mapelli M (2016) Molecular mechanisms of asymmetric divisions in mammary stem cells. EMBO Rep 17: 1700–1720

54. Stone EM, Yamano H, Kinoshita N, Yanagida M (1993) Mitotic regulation of protein phosphatases by the fission yeast sds22 protein. Curr Biol 3: 13–26

55. Tzur YB, Egydio de Carvalho C, Nadarajan S, Van Bostelen I, Gu Y, Chu DS, Cheeseman IM, Colaiacovo MP (2012) LAB-1 targets PP1 and restricts Aurora B kinase upon entrance into meiosis to promote sister chromatid cohesion. PLoS Biol 10: e1001378

56. Vaudano AP, Schwager F, Gotta M, Barbieri S (2024) Internal feedback circuits among MEX-5, MEX-6, and PLK-1 maintain faithful patterning in the Caenorhabditis elegans embryo. Proc Natl Acad Sci U S A 121: e2407517121

57. Verbinnen I, Ferreira M, Bollen M (2017) Biogenesis and activity regulation of protein phosphatase 1. Biochem Soc Trans 45: 89–99

58. Virshup DM, Shenolikar S (2009) From promiscuity to precision: protein phosphatases get a makeover. Mol Cell 33: 537–545

59. Wurzenberger C, Held M, Lampson MA, Poser I, Hyman AA, Gerlich DW (2012) Sds22 and Repo-Man stabilize chromosome segregation by counteracting Aurora B on anaphase kinetochores. J Cell Biol 198: 173–183

60. Xin G, Fu J, Luo J, Deng Z, Jiang Q, Zhang C (2020) Aurora B regulates PP1gamma-Repo-Man interactions to maintain the chromosome condensation state. J Biol Chem 295: 14780–14788

61. Zhao P, Teng X, Tantirimudalige SN, Nishikawa M, Wohland T, Toyama Y, Motegi F (2019) Aurora-A Breaks Symmetry in Contractile Actomyosin Networks Independently of Its Role in Centrosome Maturation. Dev Cell 48: 631–645 e636

