## Supplementary material for "SDS-22 stabilizes the PP1 catalytic subunits GSP-1/-2 contributing to polarity establishment in *C. elegans* embryos": Supplememntary Information

**Li et al. Supplementary Information**

Appendix Supplementary Figures EV1-EV9

Appendix Supplementary Tables 1-8

**Li et al., Fig EV1**

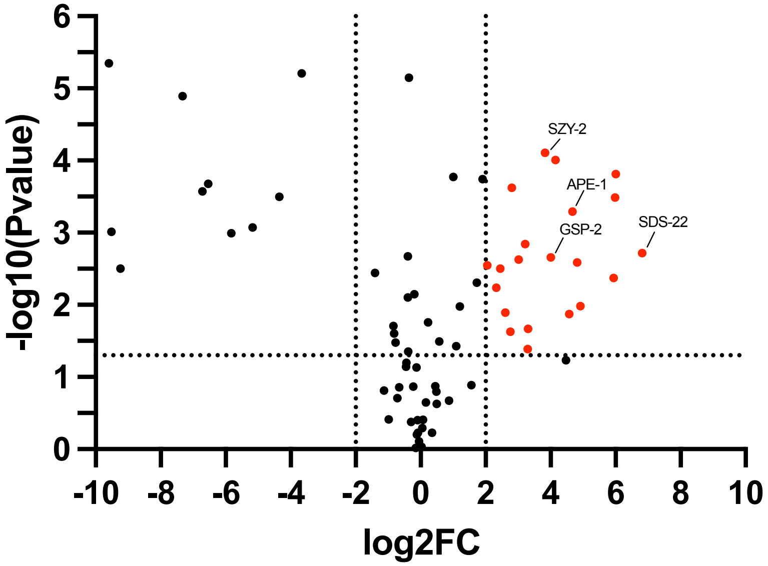


**Volcano plot of proteins identified in the GSP-2 immunoprecipitation.**
Volcano plot showing the GSP-2 interactors. The proteins highlighted are the identified regulators of GSP-2 using a threshold of log2FC≥2 (dotted lines on the log2FC≥2 axis). The Y axis is the -log10 (P value) and the dotted line represent a P value < 0.05.

### Li et al, Fig EV2

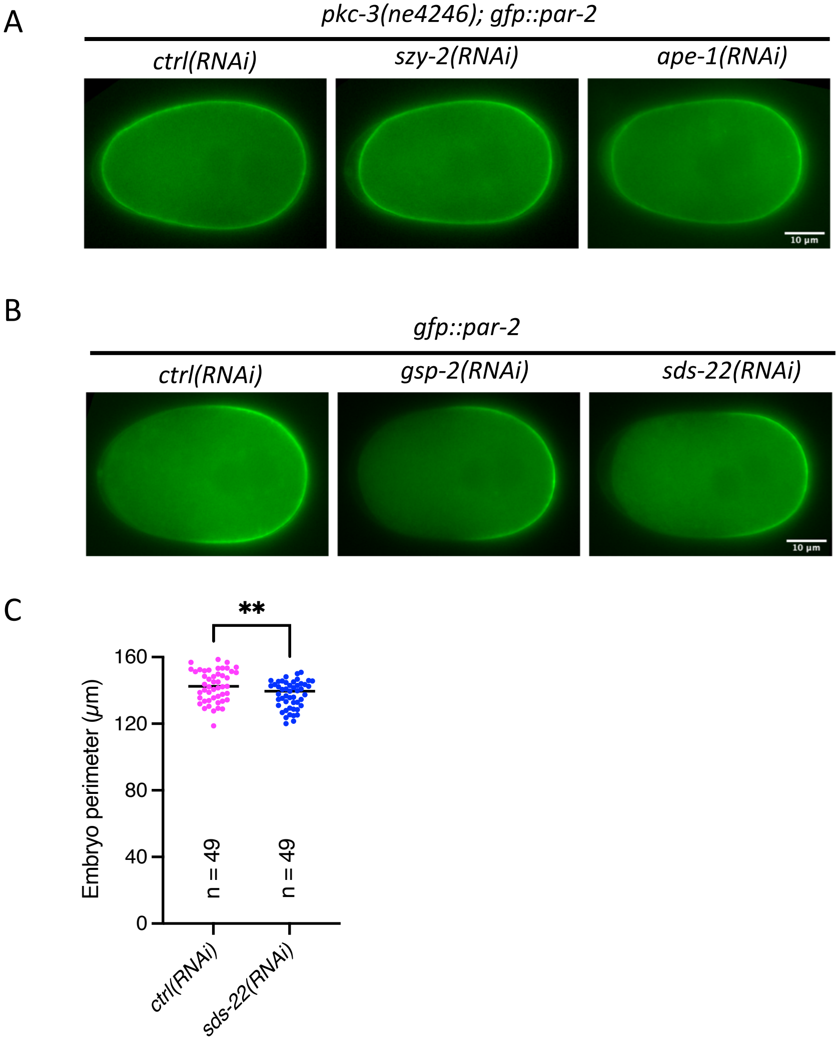


**PAR-2 localization after depletion in different genetic backgrounds of candidate regulators of PP1.**

**(A)** Still images of embryos at the pronuclear meeting stage taken from time-lapse videos: *pkc-3(ne4246); gfp::par-2; ctrl(RNAi),* n = 18, *szy-2(RNAi*), n = 17, N = 4, and *ape-1(RNAi),* n = 16, N = 2. **(B)** Still images of embryos at the pronuclear meeting stage taken from time-lapse videos of *gfp::par-2*, comparing *ctrl(RNAi),* n = 33*, gsp-2(RNAi),* n = 24 and *sds-22(RNAi),* n = 35. N = 3. The quantification of the size of the PAR-2 domain is in Fig 1C. RNA interference was performed by feeding. For all embryos, the scale bar is 10 µm, anterior is to the left and posterior to the right. n = number of embryos analyzed; N = number of independent experiments. **(C)** Quantification of embryo perimeter with indicated RNAi conditions. For both *ctrl(RNAi)* and *sds-22(RNAi)*, n = 49, N = 6.

### Li et al., Fig EV3

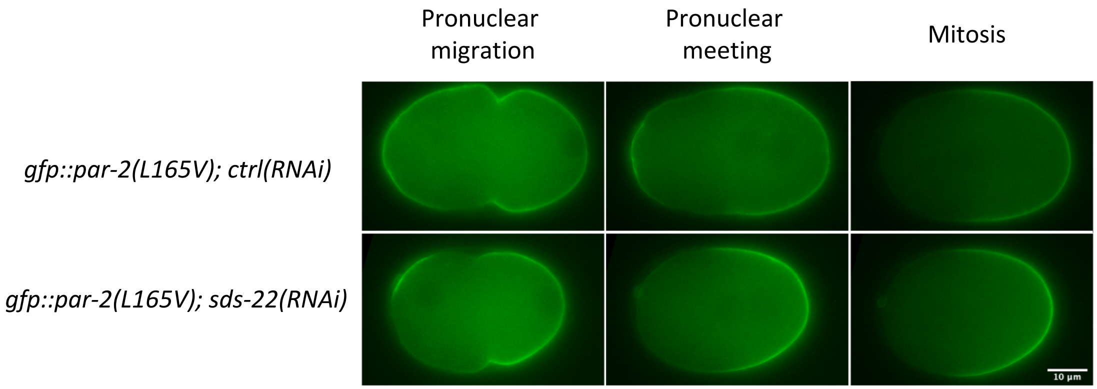


| | Pronuclear migration 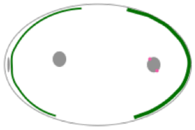 | Pronuclear meeting 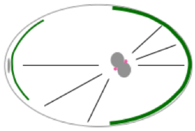 | Mitosis 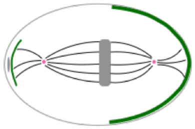 |
|---|---|---|---|
| *gfp::par-2(L165V); ctrl(RNAi)* | 100% | 100% | 14.3% |
| *gfp::par-2(L165V); sds-22(RNAi)* | 45.5% | 27.3% | 0% |

**Depletion of SDS-22 rescues aberrant PAR-2 localization in *gfp::par-2(L165V)* mutant.**

Top: still images from time lapse videos of *gfp::par-2(L165V)* of one-cell embryos at different cell division stages, *ctrl(RNAi),* n = 9 and *sds-22(RNAi)*, n = 11, N = 3. Bottom: the table show the percentage of the phenotype represented schematically at the top . RNA interference was performed by feeding.

### Li et al., Fig EV4

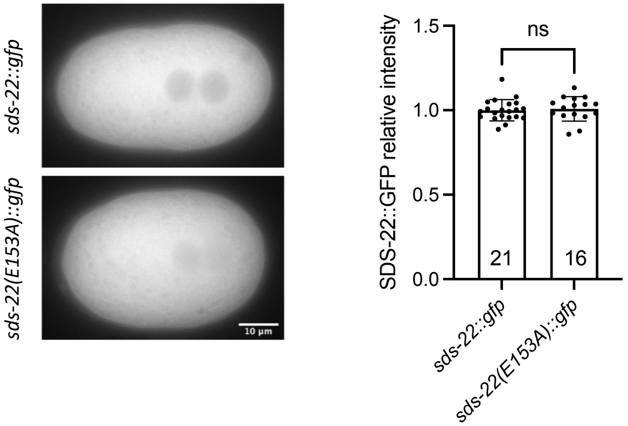


**The E153A substitution does not result in reduced levels of SDS-22.**

Images of *sds-22::gfp* and *sds-22(E153A)::gfp* at pronuclear meeting stage, left and quantification of relative levels of SDS-22(E153A)::GFP normalized to SDS-22::GFP levels, right. Mean is shown and error bars indicate SD. Each dot represents a single embryo. Sample size (n) is indicated inside the bars in the graph. N = 2. ns $p > 0.05$. The P-values were determined using two-tailed unpaired Student's t-test.

### Li et al., Fig EV5

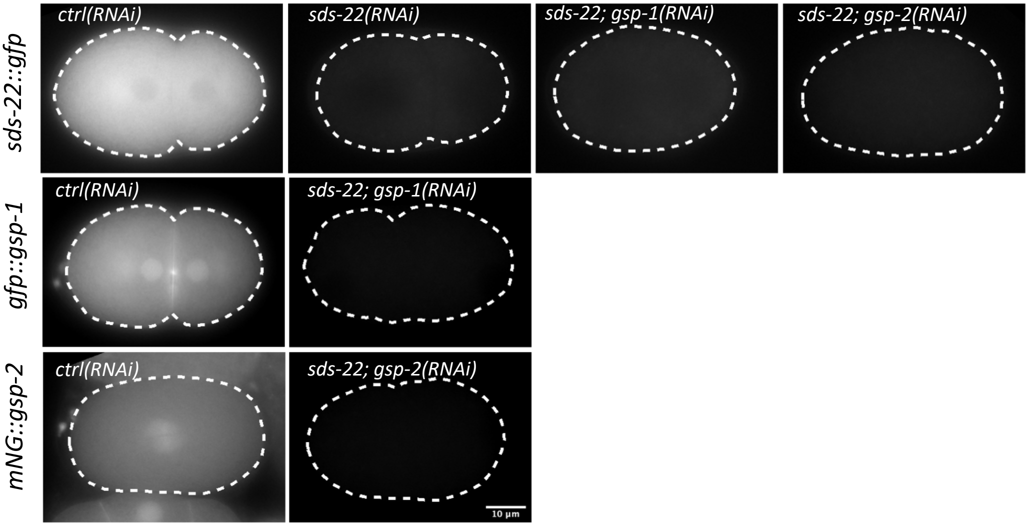


**SDS-22, GSP-1 and GSP-2 were depleted in co-depletion experiments.**
Representative images of the indicated genotypes. Co-depletion of SDS-22 with GSP-1 or GSP-2 is efficient. RNA interference was performed by injection.

**Li et al., Fig EV6**

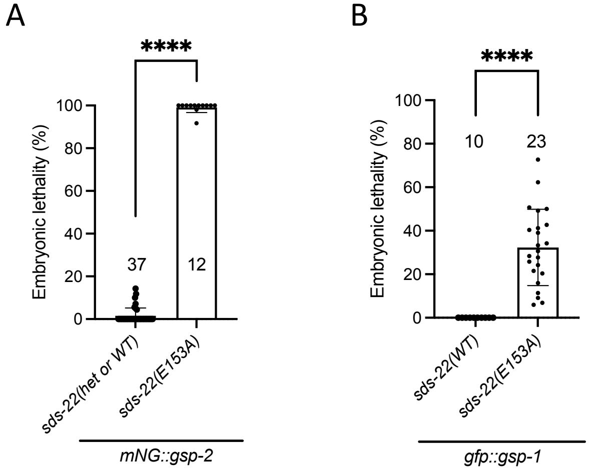


**The E153A substitution in SDS-22 leads to embryonic lethality in the *mNG::gsp-2* and *gfp::gsp-1* genetic backgrounds.**

**(A, B)** Embryonic lethality of embryos with the *sds-22(E153A)* mutation in the genetic background of *mNG::gsp-2* **(A)** and *gfp::gsp-1* **(B)**. The reported values correspond to the percentage of un-hatched embryos over the total progeny (larvae and un-hatched embryo). *mNG::gsp-2; sds-22(E153A/+ or +/+),* n = 1703, *mNG::gsp-2; sds-22(E153A),* n = 568. N = 3. *gfp::gsp-1,* n = 713, *gfp::gsp-1; sds-22(E153A),* n = 1783. N = 2. In this plot, each dot represents for the quantified lethality of one single plate. The P-values were determined using two-tailed unpaired Student's t-test. ****p < 0.0001.

### Li et al., Fig EV7

A

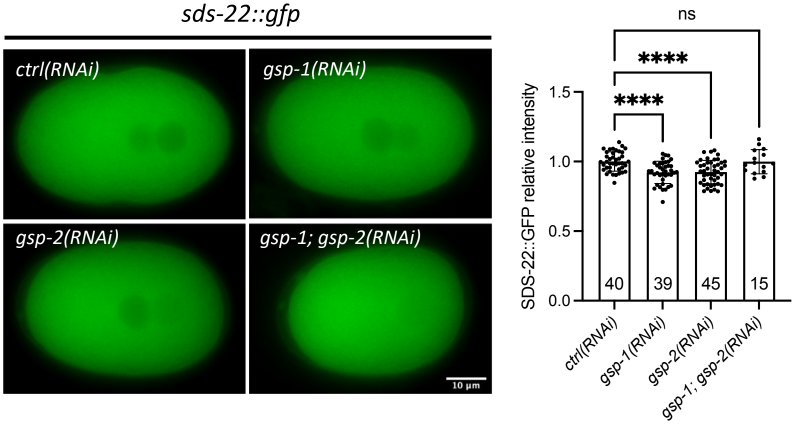


B

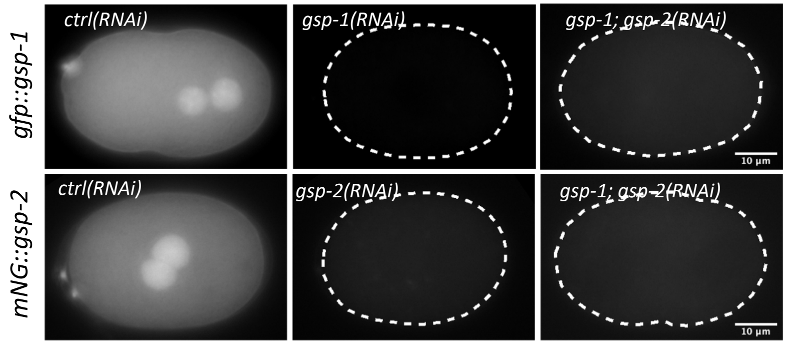


**SDS-22 levels are not affected by depletion of GSP-1 and/or GSP-2.**

**(A)** Left, representative images of *sds-22::gfp* embryos in *ctrl(RNAi), gsp-1(RNAi), gsp-2(RNAi)* and *gsp-1(RNAi); gsp-2(RNAi)*. Right, quantification of relative SDS-22::GFP levels. Mean is shown and error bars indicate SD. Sample size (n) is indicated inside the bars in the graph. N = 3. ns $p > 0.05$, ****$p < 0.0001$. The P-values were determined using one-way ANOVA "Tukey's multiple comparisons test". **(B)** Representative images of the indicated genotypes. GSP-1 and GSP-2 are efficiently depleted in the indicated co-depletions. RNA interference was performed by injection.

### Li et al., Fig EV8

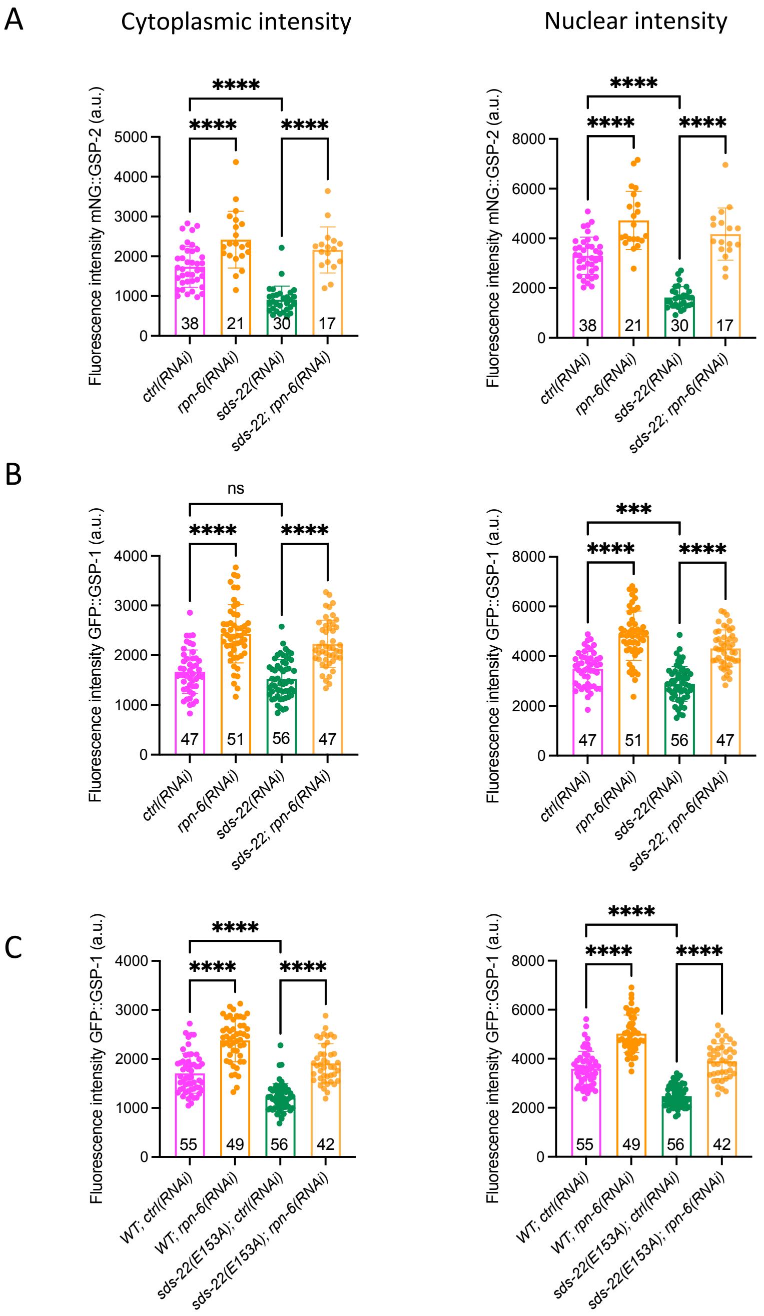

**Depletion of RPN-6.1 shows the same effect as RPN-7 depletion.**
**(A-B)** Quantification of mNG::GSP-2 **(A)** and GFP::GSP-1 **(B)** intensity levels in the cytoplasm and nucleus of -1 and -2 oocytes in the *ctrl(RNAi), sds-22(RNAi), rpn-6(RNAi),* and *sds-22(RNAi); rpn-6(RNAi)* conditions. Mean is shown and error bars indicate SD. The P-values were determined using one-way ANOVA "Tukey's multiple comparisons test". Sample size (n) is indicated inside the bars in the graph. N = 3. **(C)** Quantification of GFP::GSP-1 intensity levels in the cytoplasm and nucleus of -1 and -2 oocytes, comparing wild type, SDS-22(E153A) mutation, *ctrl(RNAi)* and *rpn-6(RNAi)*. Mean is shown and error bars indicate SD. The P-values were determined using one-way ANOVA "Tukey's multiple comparisons test". Sample size (n) is indicated inside the bars in the graph. N = 3. The RNP-6.1 has been performed the same days as the RNP-7 depletion assay of Fig. 7, and it is therefore the same *ctrl(RNAi)* and *sds-22(RNAi)* data as shown in Fig. 7B, C, E, F, H and I. For all panels, RNA interference was performed by feeding. The scale bars are 50 µm. In all plots, ns $p > 0.05$, * $p < 0.05$, ** $p < 0.01$, ***$p < 0.001$, ****$p < 0.0001$, n = number of embryos analyzed; N = number of independent experiments.

#### Li et al., Fig EV9

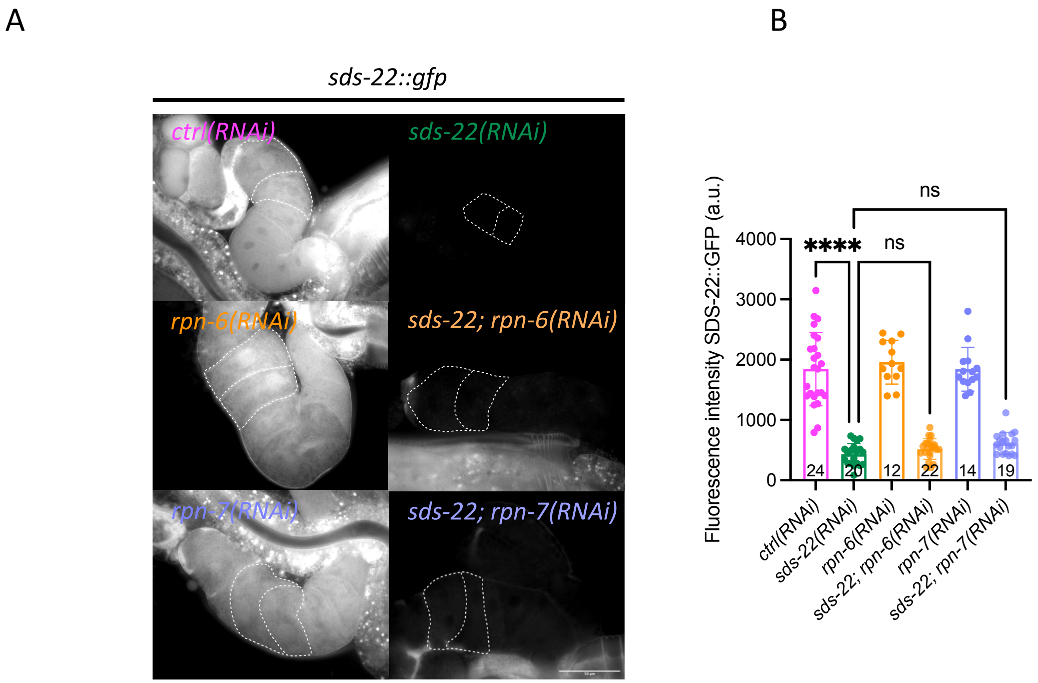


**Depletion of SDS-22::GFP is efficient in the co-depletion experiments.**
**(A)** Representative midsection images of *sds-22::gfp* germlines in *ctrl(RNAi), sds-22(RNAi), rpn-6(RNAi)* or *rpn-7(RNAi)* single depletion, and *sds-22(RNAi); rpn-6(RNAi)* and *sds-22(RNAi); rpn-7(RNAi)* co-depletion. No viable zygotes were produced after depletion of RPN-6/-7 alone or co-depletion with SDS-22, indicating that RPN-6 and RPN-7 were well depleted in these conditions. Scale bar is 50 µm. **(B)** Quantification of SDS-22::GFP intensity levels in the cytoplasm of -1 and -2 oocytes. Mean is shown and error bars indicate SD. The P-values were determined using one-way ANOVA "Tukey's multiple comparisons test". Sample size (n) is indicated inside the bars in the graph. N = 3. ns $p > 0.05$, ****$p < 0.0001$.
